# Super-Resolution Imaging of Fast Morphological Dynamics of Neurons in Behaving Animals

**DOI:** 10.1101/2023.10.23.563526

**Authors:** Yujie Zhang, Lu Bai, Xin Wang, Yuchen Zhao, Tianlei Zhang, Lichen Ye, Xufei Du, Zhe Zhang, Jiulin Du, Kai Wang

## Abstract

Neurons are best studied in their native working states in which their functional and morphological dynamics support animals’ natural behaviors. Super-resolution microscopy can potentially reveal these dynamics in much higher details but has been challenging in behaving animals due to severe motion artifacts. Here, we report multiplexed line-scanning structured illumination microscopy, which could tolerate motions of up to 50 μm/s while achieving 150 and 100 nm lateral resolutions in its linear and nonlinear forms respectively over large fields-of-view. We continuously imaged the dynamics of spinules in dendritic spines and axonal boutons volumetrically over thousands of frames and tens of minutes in mouse brains during sleep-wake cycles. Super-resolution imaging of axonal boutons revealed their prevalent spinule dynamics on a scale of seconds. Simultaneous two-color imaging further enabled analyses of the correlation between the spatial distributions of diverse PSD-95 clusters and the structural dynamics of dendrites in the brains of awake mice.

## INTRODUCTION

Neurons are highly specialized in morphology to support their complex functions in information integration, transformation, and communications^1,2^. In particular, these morphological specializations, such as dendritic spines, axonal boutons and synaptic connections between them, undergo constant changes throughout an animal’s lifespan to enable behavioral adaptations that are critical for the animal’s survival in varying environments^3–5^. Imaging these morphological dynamics in behaving animals at higher spatiotemporal resolution can drive better understanding of the physiology of neurons and their networking into functional and plastic circuits.

Morphological studies of neurons are traditionally performed using electron microscopy (EM) on fixed tissues^6^. However, such a single time-point observation approach has low efficiency in studying dynamic changes in highly complex and plastic neural circuits. In comparison, fluorescence microscopy allows longitudinal investigation of the same neurons over time in behaving animals. In particular, two-photon microscopy has been widely applied in the structural imaging of neurons and advanced our understanding of structural neural plasticity^3,7^. However, many critical details of neurons are beyond the resolving capabilities of conventional fluorescence microscopy and require correlative EM dissection following optical imaging^8^. To bridge this resolution gap, super-resolution microscopy (SRM) has been applied for optical neural imaging and proven to be valuable in studying fine structures and molecular organizations at nanoscales^9–14^.

SRMs can overcome the resolution limit (∼250 nm) imposed by diffraction, but they usually have much reduced imaging speed and have been mainly applied in imaging cultured cells and tissue sections^9,12,15–18^. Achieving super-resolution imaging in living animals has remained a challenging frontier because any minute motion inevitable in a living animal can lead to significant imaging artifacts. To alleviate this problem, animals were anesthetized in recent demonstrations of stimulated-emission-depletion (STED) microscopy imaging of mouse brains^10,19–22^. Notably, morphological features and longitudinal changes in them can be tracked at resolutions below 100 nm. Structured illumination microscopy (SIM) has also been applied to image anesthetized mouse brains where sample motions were computationally corrected^23^. Anesthetization, however, significantly alters the normal physiology of neural circuits and animals’ behaviors and the field is still missing a super-resolution technique that is applicable in awake and behaving animals.

Here, we introduce multiplexed line-scanning structured illumination microscopy (MLS-SIM), a new type of SRM that enables longitudinal super-resolution imaging in brains of awake and behaving animals, is free of motion-induced artifacts, and be performed on large fields of view. MLS-SIM can tolerate severe sample motion of up to ∼50 µm/s by employing a line-scanning imaging scheme. It achieves super-resolution imaging in all three dimensions within a single continuous scan of multiplexed and patterned line excitations. In this way, any nanoscale structure spanning a few adjacent imaging lines can be captured within several hundreds of microseconds and resolved at super-resolution without being influenced by the often severe sample motions. We demonstrated the capabilities and robustness of MLS-SIM by imaging and tracking morphological dynamics and molecular organizations in dendritic spines and axonal boutons over hundreds of time points during the awake and sleep states of mice. By making use of fluorescence saturation effect^24,25^, the nonlinear version of MLS-SIM was able to further enhance the spatial resolution without compromising the tolerance for sample movement.

## RESULTS

### Principles of MLS-SIM

SIM is one of major types of SRMs that breaks the resolution limit of fluorescence microscopy by applying non-uniform excitation patterns, including sinusoidal gratings^26^, structured lines^27^, diffraction-limited focused spots^28,29^, and even random patterns^30^. Among these patterns, structured line is the best candidate for imaging awake animals due to its two critical features: effective background rejection by confocal line detection and high-speed imaging by parallelized imaging along the line. However, previous attempts required multiple scans over the same field of view to reconstruct one super-resolved image^27,31^ and thus could not tolerate sample motions well in awake animals. To address this problem, we developed MLS-SIM that can super-resolve in three directions after a single scan cycle with only one readout required for each line, and can therefore super-resolve local structures even in the presence of severe global sample motions.

MLS-SIM was built on the platform of a confocal line-scanning microscope (Figures 1A and S1A, and STAR Methods). To improve the resolution in all three directions, two structured line illumination patterns were applied. One is the traditional focused line with constant intensity along the x direction and tightly focused to the diffraction limit in the y direction (Figures 1A and 1B). The other pattern introduced periodic intensity modulations along both the x and the z directions (Figures 1A and 1B), similar to that used in the structured line illumination microscopy^27^ except that ours used three-beam, instead of two-beam, interference in a manner similar to 3D-SIM^26^. The sharpness of the focused line and the interference spots are critical to the final resolution gain. To ensure that, the laser beams were polarized along the x and the y directions for two different patterns, respectively (Figure 1B). For fluorescence detection at each scan step, a de-scanned 2D image was collected by a slim region of interest on the camera centered along the excitation line (Figure 1A). During image acquisition, three operations were performed simultaneously (Figures 1A and S1B): (1) Excitation lines were scanned in the y direction to cover a 2D field of view; (2) The two excitation patterns were alternated; and (3) The phase of the second and periodic pattern was advanced in the x direction.

**Figure 1.**
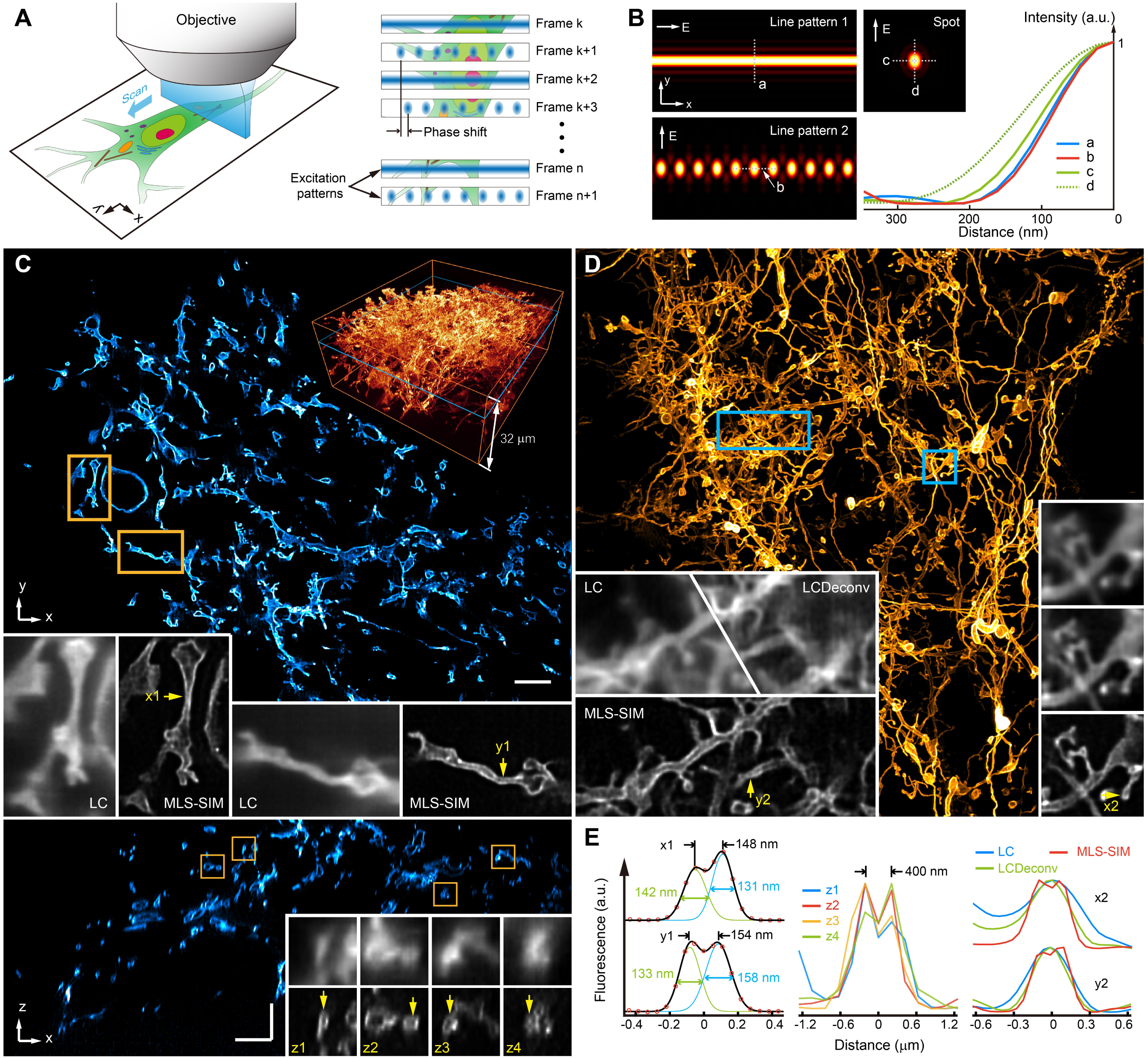
Multiplexed-line-scanning structured-illumination-microscopy (MLS-SIM). (A) The concept of spatial and temporal multiplexing of two different excitation line patterns in MLS-SIM. The line scans in the y direction. (B) Simulated excitation line patterns and their comparisons with a conventional focused spot. The linear polarizations used for generating the excitation patterns are indicated by arrows denoted by “E”. The comparison of intensity line profiles at indicated locations shows tighter light focusing in the y and the x directions than a conventional focused spot. (C) Volumetric imaging of Purkinje cells expressing membrane-targeted EGFP by MLS-SIM in living larval zebrafish. Representative x-y view (top) and x-z view (bottom) of single planes in the volume. 3D rendering of the volume is shown at the top-right. Bottom insets compare resolution between conventional line scan confocal and MLS-SIM in magnified regions as indicated. (D) Maximum Intensity Projection (MIP) of an imaging volume (20 planes covering 4 µm in the z direction) of neurons expressing membrane-targeted EGFP in an awake mouse brain. Insets are magnified views of selected regions at single planes in the volume to highlight super-resolution by MLS-SIM. (E) Intensity line profiles at locations indicated by arrows in (C) and (D). Line profiles were fitted by Gaussian functions to estimate the sizes of resolvable features. The full width at half maximums (FWHM) of the fitted Gaussian functions and their separations are specified. LC, raw line confocal imaging; LCDeconv, line confocal imaging with deconvolution. Scale bars: 5 µm in (C).

When acquiring an image, each pixel on the camera can be considered as a photodetector of a separate confocal microscope and detected with its own effective point spread function (PSF), which is the multiplication between the excitation pattern and detection PSF. Different photodetectors exhibit different effective PSFs depending on their spatial interrelationship with the excitation pattern (Figures S2A and S2B). These effective PSFs had sharper spatial profiles than the PSFs of the confocal microscopy and served as the basis for the super-resolution in MLS-SIM. The first illumination line pattern improved the resolution in the y direction, while the second improved the resolution in the x and the z directions (Figures 1B and S1E). Multiplexing these two patterns with a single scan, in principle, captured complementary high-resolution information in all three directions, however, how to properly extract these information from the raw image strips and coherently combine them into one super-resolution reconstruction was very challenging and required a completely new algorithm. Due to the pattern alternation and the phase scanning of the second pattern, the same camera pixel saw different excitation pattern, resulting in different effective PSFs, at different scan steps. To resolve this complication, our algorithm first divided the pixels of all raw image strips from different scan steps into N different groups (Figure S2) based on the type of effective PSFs governing each pixel. It then assigned or duplicated each raw-image pixel, based on which group it belongs to, to one of N images stitched row-by-row from the line scans, one image for each group. Each pixel’s row assignment in a stitched image corresponds to the scan step from which it originated. Each of these N images was considered to have been acquired under its own unique effective PSFs and they are then jointly deconvolved to reconstruct the final super-resolved image. In addition, this reconstruction framework could make full use of 3D PSFs to perform multi-layer reconstructions on 2D images to offer super-resolution optical sectioning in the z direction (STAR Methods and Figure S4).

Compared with wide-field super-resolution techniques such as localization microscopy^16,17^ and SIM^26^, MLS-SIM does not need to accumulate multiple frames of the same sample area to obtain one super-resolution image and therefore, can better tolerate sample motion and capture local structures at super-resolution within hundreds of microseconds. Compared with spot-scanning based super-resolution imaging techniques, MLS-SIM uses much lower peak laser intensity, resulting in a lower photobleaching rate and reduced phototoxicities, and can produce a larger field of view more easily because of one less dimension to scan.

### MLS-SIM enabled super-resolution imaging in awake animals

We first characterized the resolution of MLS-SIM by imaging Purkinje cells expressing membrane-targeted EGFP in larval zebrafish. Due to the confocal line detection, interfering background was effectively rejected and volumetric imaging could be performed on densely labeled samples (Figure 1C and Video S1). The imaging depth was limited by tissue-induced aberrations and scattering^23^. Examining the smallest features that could be resolved laterally in an exemplar imaging plane about 15 µm below the sample surface demonstrated a practically achievable lateral resolution of around 150 nm (Figure 1E). In the axial direction, MLS-SIM achieved optical sectioning and super-resolution of 400 nm (Figure 1E). Unlike wide-field 3D-SIM that requires 3D stack acquisition, MLS-SIM only requires single plane acquisition and this feature makes the latter more tolerant to motion-related distortion when imaging an awake animal. To highlight the benefit of this feature, we imaged neurons expressing membrane-targeted EGFP in awake mice brains (Figure 1D and Video S2). Even though the image was distorted overall by sample motion, super-resolution was still preserved in imaging local structures. Furthermore, the super-resolution optical sectioning capability of MLS-SIM greatly enhanced imaging contrast for in-focus features embedded in densely labeled regions in the surrounding planes (Figure 1D).

To quantify the tolerance of sample motions in MLS-SIM imaging, we imaged fluorescent particles intentionally moved at three different speeds (25, 50, and 100 µm/s) and in eight different directions (Figures 2A, S3A and S3B). We identified pairs of fluorescent beads that could be resolved under different sample motions and estimated achievable resolution limits in different directions by fitting curves to these data points (STAR Methods). When the sample was kept static during imaging, the estimated resolution was 120 nm and 150 nm in the x and y directions, respectively. These resolutions were well preserved when samples were moved at a speed of 25µm/s. As the speed was increased to 50 µm/s, the resolutions could still be preserved except for a few directions in which the sample was moved. For example, the resolution along the y direction was most severely degraded to ∼200 nm when the sample was moved in that direction. When averaging over all motion directions, the estimated resolutions along the x and y directions were kept below 150 nm. When the speed was further increased to 100 µm/s, the resolutions along the directions perpendicular to the moving directions could still be preserved (Figure S3C), even though reconstructed images were severely distorted.

Even though MLS-SIM only needs a single plane to achieve super-resolution sectioning, single-plane imaging would not be able to track axial movements that are inevitable during longitudinal studies. We therefore performed volumetric imaging in all longitudinal studies to allow tracking and at each time point, corrected the motion-induced image distortions using a row-by-row image registration pipeline (Figure 2B and STAR Methods). In one such experiment in a head-fixed mouse running on a wheel, the average speed of brain motion was found to be 23 µm/s. More than 7.6% of the time, the sample moved at speeds higher than 50 µm/s (Figure 2C). Nonetheless, the super-resolution imaging performance of MLS-SIM was shown to be well preserved, even in the presence of instantaneous movements at a speed exceeding 100 µm/s (Figures 2D and 2E, Video S3).

**Figure 2.**
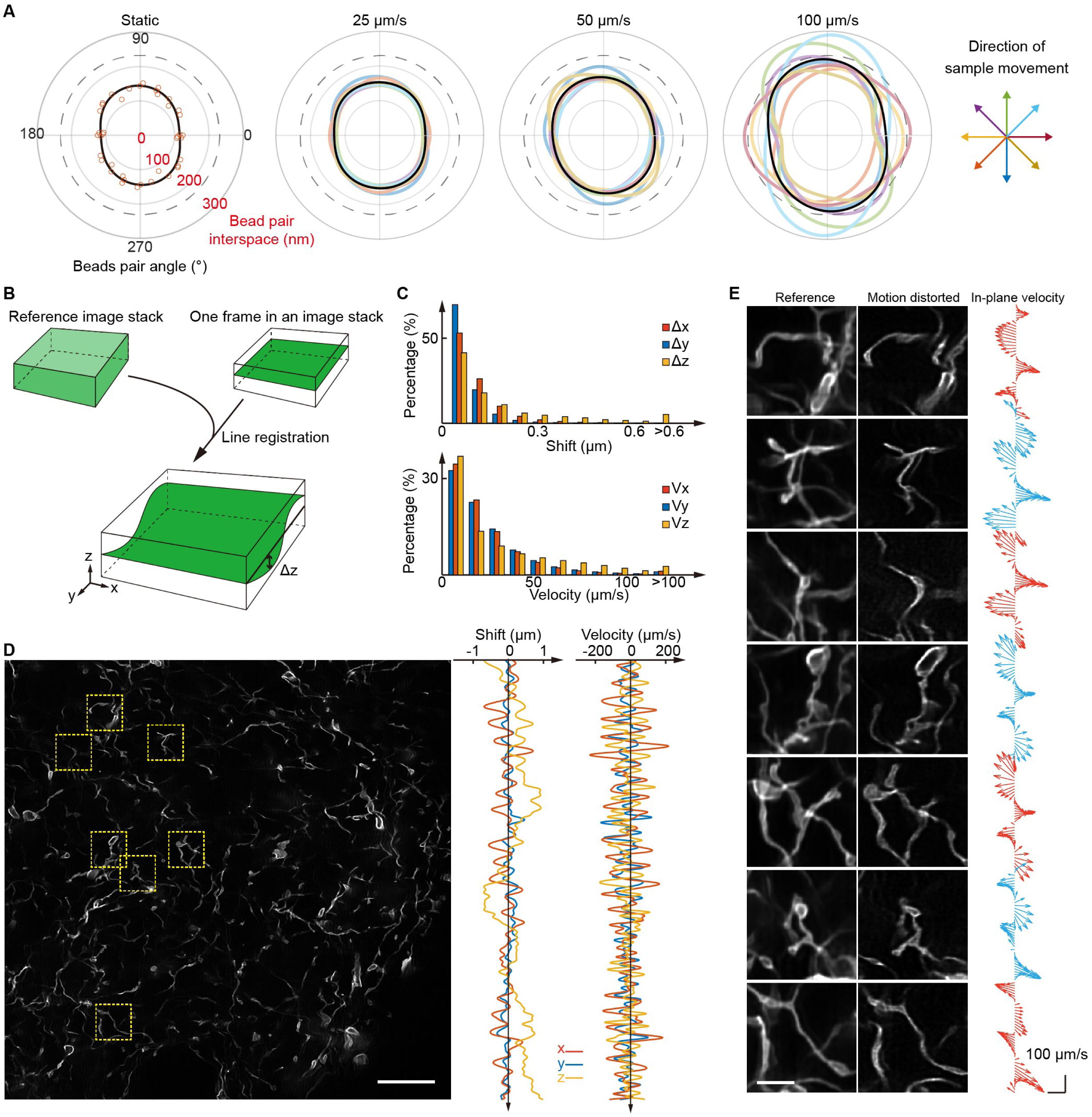
Motion tolerance of MLS-SIM and correction of motion induced image distortions. (A) Experimental characterization of lateral resolutions in the presence of sample motions by imaging 100 nm diameter fluorescent beads. Intentional sample motions were introduced in eight directions, indicated by different colors, and at three different speeds. Colored circles were fitted to experimentally measured minimal interspaces on resolvable bead pairs (Figures S2A and S2B, STAR Methods) and the black circles represent the average of colored circles measured over the eight motion directions. The dashed gray circles indicate the theoretical diffraction limit of the system. (B) Schematics of the image registration algorithm. Each motion-distorted image stack is aligned with respect to a reference image stack, which is typically constructed out of a time series of image stacks (STAR Methods). Each row in a captured frame can suffer from severe motions in the axial direction and its motion-induced shift can be estimated by performing an exhaustive search of the best match in the reference image stack. (C) Distributions of spatial shifts and instantaneous velocities of sample motions in an awake mouse brain estimated from the row-by-row registration. (D) A representative motion-distorted imaging frame captured in an awake mouse brain. The spatial shifts and instantaneous velocities estimated for each row in this frame are plotted on the right. Motion speed could occasionally exceed 200 µm/s. (E) Magnified images of the indicated ROIs in (D) and their comparisons with the same ROIs in the reference image stack. The instantaneous velocities in the x-y plane are plotted as vectors on the right. High instantaneous velocity of sample motion markedly distorted images. However, small features could still be resolved at super-resolution most of the time. Scale bars: 10 μm in (D); 2 μm in (E).

### Tracking fast morphological dynamics of neurons at super-resolution in awake mice brains

The combination of MLS-SIM and image registration pipeline offers a unique opportunity to investigate the morphological dynamics of neurons at super-resolution in awake mice brains. The morphological specializations of neurons, including dendritic spines and axonal boutons, have been studied extensively due to their essential role in neural circuitry^32,33^. In particular, in vivo studies using two-photon microscopes showed that the dynamic changes of spines are responsive to animals’ experiences and environmental changes^3,7^. However, spines are highly diverse in structures and dynamics^32^ and require morphological characterizations at higher resolutions to better investigate their correlations with various forms of functional and behavioral changes in awake and behaving animals. For example, many dendritic spines have spinules, which are thin protrusions originating on spine heads and first detected by EM^34^. Spinules are found widespread in different brain areas^35^, but they can hardly be detected in vivo using two-photon microscopes due to their small sizes. To study their dynamics, SRMs were recently introduced to image spinules on cultured neurons and brain slices^36^. Further super-resolution studies of their dynamics under natural physiological conditions and animals’ behavioral states can greatly advance our understanding of their in vivo functions.

We therefore labeled excitatory neurons by membrane-localized EGFP in mouse primary somatosensory cortex and imaged dendritic trees of the labeled neurons in cortex layer I through a chronic glass window. Since spines and spinules undergo morphological changes with a time scale of tens of seconds, we chose to image a volume of 80 × 40 × 4 μm^3^ (1536 × 768 × 20 voxels) with a temporal resolution of 8 seconds and performed continuous time-lapse imaging with MLS-SIM over tens of minutes. With optimal fluorescent labeling in one example, we were able to continuously image a volume of 80 × 20 × 4 μm^3^ over 298 time points, containing 5960 frames, in 20 minutes (Figure S3 and Video S4). Except for a few time points when occasional extreme movements were uncorrectable, the same structure could be tracked over time readily after image registration. Spinules could be identified as dynamic protrusions on various types of spines, ranging from large mushroom-shaped to thin spines (Figure 3A and Video S5). By analyzing 457 spines imaged in 14 volumes in 5 mice, we found 62% (285 in 457) of them exhibited spinule dynamics (1419 spinules) within the recording time period (Figure 3B), which was similar to previous findings from studies on ex vivo brain slices^36^. Most spinules were small in sizes and emerged highly transiently (Figure 3C). A few spinules could extend more than one micrometer from the spines’ heads (Figure 3D). The exploration ranges of spinules were positively correlated with their lifetimes (Figure 3C). Some spines exhibited a higher level of spinule dynamics by having spinules emerging from multiple sites on the same spine or containing “active sites” that generated spinules repeatedly (Figures 3E and 3F). We found 68% (194 in 285) of spines that had spinules contained two or more sites for spinule emergence, and 46% of all these sites (323 in 706) were “active” in having more than one protrusion-retraction cycle of spinules. In one example, we observed spinule recurrence of 14 times within 28 minutes (Figure 3E), indicating their high level of dynamics in vivo.

**Figure 3.**
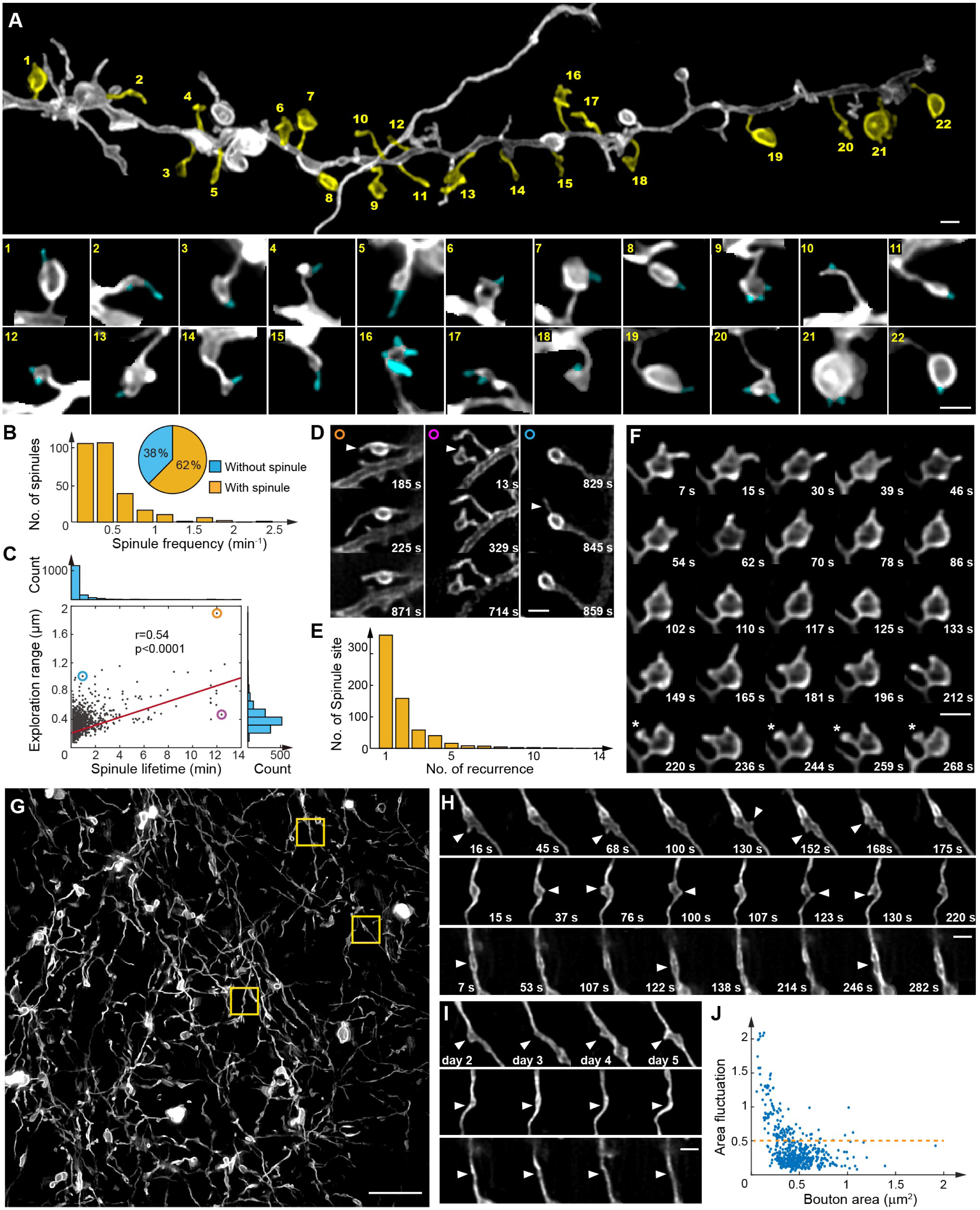
Imaging fast morphological dynamics of neurons with MLS-SIM in awake mice brains. (A) Overview (top) of a dendrite branch and magnified views (bottom) of indicated dendritic spines, as numbered in the overview, captured at particular time points when spinule dynamics appeared. The identified spinules were highlighted in cyan. (B) Histogram of spinule emergence frequencies. The pie chart shows the percentage of spines experiencing spinule emergences. (C) Dependence of exploration range on the lifetime of the observed spinules. The exploration range was found to be positively correlated with the spinule’s lifetime (r=0.54, p<0.0001, Pearson’s correlation). (D) Images of selected spinules, corresponding to the circled data points in (C), with different lifetimes and exploration ranges. (E) Histogram of the number of spinule sites that exhibited recurrent emergences of spinules. (F) A representative spine that had multiple spinule emergence sites and recurring spinules. The asterisks indicate a mushroom-spine-shaped spinule. (G-I) Overview of axons of somatostatin interneurons expressing membrane-targeted EGFP. Magnified and time-lapsed views of indicated regions in (G) are shown in (H) for the first day and in (I) for across 5 consecutive days. (J) Distributions of dynamic area changes over boutons (n=535). The area fluctuation is defined as the standard deviation of a bouton’s area change normalized to its average area across 5 days. Scale bars: 1 µm in (A), (D), (F), (H) and (I); 10 µm in (G).

Compared to dynamic dendritic spines, excitatory axonal boutons in the cortex are relatively stationary, as suggested by in vivo two-photon imaging studies on mice^37^. However, EM studies reported that spinules could also be formed on presynaptic boutons, more typically on inhibitory ones^35^. These bouton spinules, known as pseudopodial indentation^35^, have yet to be studied for their dynamics by light microscopy. We labeled axons of somatostatin positive interneurons by membrane-localized EGFP in the primary somatosensory cortex in mice and imaged with MLS-SIM a volume of 80 × 80 × 4 μm^3^ for 38 time points in five minutes every day over five successive days (Figure 3G). The results revealed small protrusions on some of the boutons (Figure 3H) similar in shape to dendritic spinules. These small protrusions also exhibited fast dynamics on the time scale of seconds (Figure 3H) and were presumably bouton spinules reported in EM studies^35^. In 535 analyzed boutons, 47% showed emergences of spinules within the imaging window. More extensive morphological changes of boutons could be observed over a longer time scale of days (Figure 3I). Complete disappearance of boutons was rare (9 in 535 cases) and 80% (427 in 535) of the observed boutons had size fluctuations below 50% of their average sizes over five days (Figure 3J).

### Imaging structural dynamics of neurons during sleep-wake cycles in mice brains

The morphological dynamics of neurons are widely believed to contribute to processes of memory and neural plasticity^3,32,33^, but it is unclear whether these dynamics are definitively different between awake and sleep states even though sleep is hypothesized to be specifically involved in certain stages of memory processes^38,39^. Pioneering studies by EM and in vivo two-photon imaging indeed reported such differences in synaptic connections^40^ and turnover rate of dendritic spines^41–43^, respectively. We set out to study these potential differences using MLS-SIM.

To maintain the natural posture of the mice during sleep, the imaging objective was positioned at an angle to keep the mouse head staying up-right during imaging experiments (Figure 4A). The sleep-wake states of mice were monitored by electroencephalogram (EEG) and electromyogram (EMG) recordings (Figure 4B). To reduce perturbation and maximize temporal coverage, we imaged at two volumes per minute. In one experiment, multiple sleep-wake cycles, comprising both non-rapid eye movement (NREM) and rapid eye movement (REM) sleep states, were captured by intermittently imaging at 51 time points in 30 minutes (Figures 4A, 4B and Video S6). The dynamic changes of 248 neural structures in one field of view, including axonal boutons, dendritic spines, and dendrite trunks with emerging filopodia, were analyzed. We observed dynamic emergences of spinules on boutons and spines. Based on this limited data, the change in the number of spinules did not show obvious correlation with sleep-wake cycles (Figure 4C). We further analyzed morphological changes of neural structures between two consecutive time points, which can be complicated because protrusions and retractions could occur simultaneously on different locations of the same neural structure (Figure 4D). We quantified these morphological changes and found that they were highly dynamic during both wake and sleep states (Figure 4E). Nevertheless, calculations of area changes used in our morphological analysis greatly simplifies the observed dynamics. More sophisticated analysis^44^ is still needed to better understand these changes and their correlations with different brain states.

**Figure 4.**
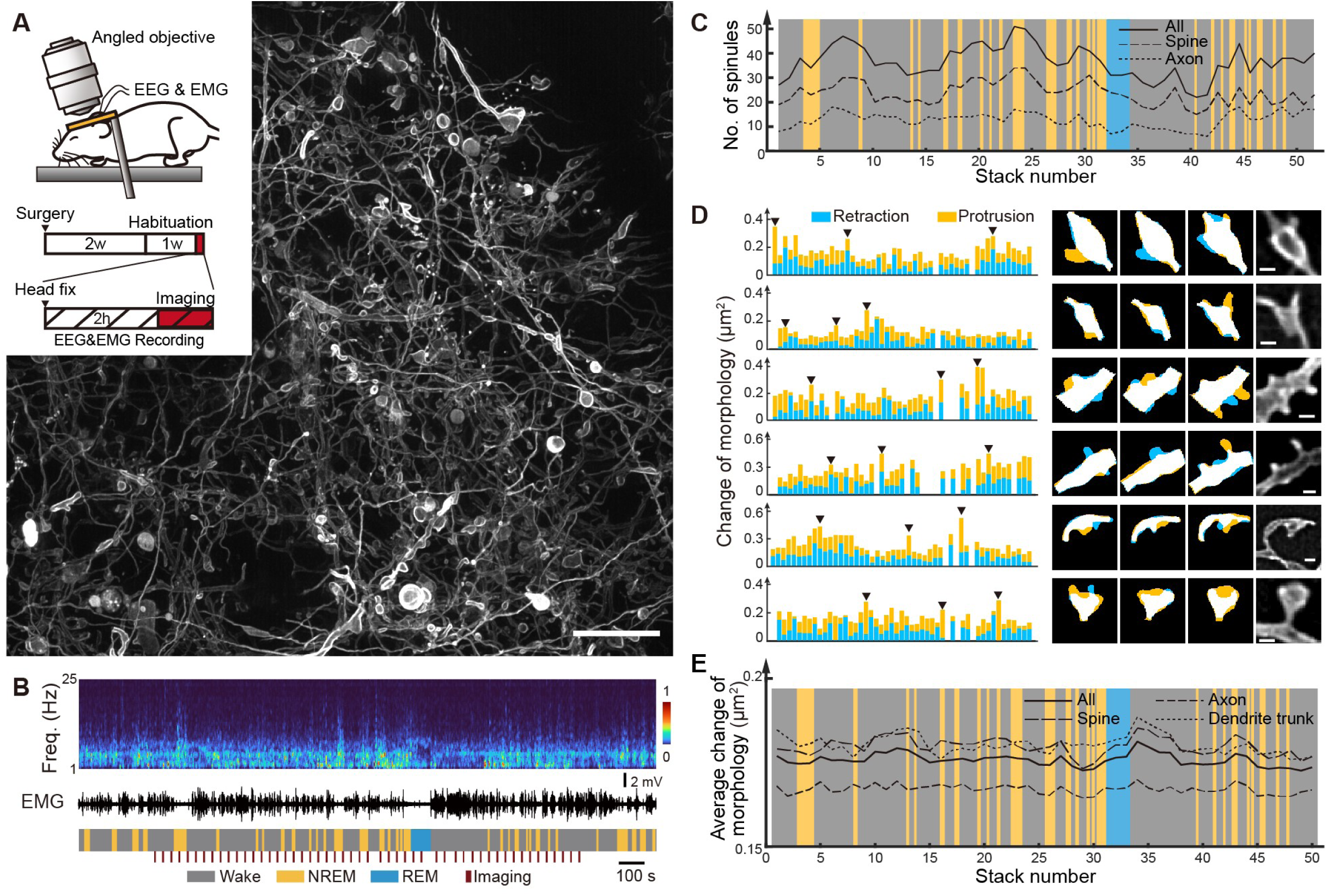
Tracking morphological dynamics of neurons with MLS-SIM during sleep-wake cycles in mice brains. (A) Overview of the imaging field captured during sleep-wake cycles. Inset: schematics of imaging mouse brain with an angled imaging objective and special experimental procedure to preserve normal sleep behavior, where the red regions indicate the time interval when volumetric MLS-SIM imaging were performed. (B) EEG spectrogram, EMG recording and sleep-wake states of the mouse. Red bars indicate the time points when volumetric MLS-SIM imaging were performed. (C) Change of the total number of spinules identified both on dendritic spines and axonal boutons during the sleep-wake cycles, indicated in the background of the graph. (D) Examples of quantifications of morphological changes of neural structures. Protrusions and retractions were quantified separately and summed to represent the total change in morphology. (E) Averaged change of morphologies as quantified in (D) during sleep-wake cycles, indicated in the background. Scale bars: 10 µm in (A); 500 nm in (D).

### Two color super-resolution imaging in awake mice brains

Although characterizing neural morphologies can provide insights on the synaptic connections, imaging of synapse-related molecules are usually needed to better examine the existence, strength, and stability of synapses^32^. For example, excitatory synaptic scaffold protein PSD-95 is widely accepted as a marker for synaptic maturation and spine stabilization and has been under extensive imaging investigations^32^. In particular, STED has been used to study their morphological diversities in anesthetized mice brains, revealing the dynamic reorganizations over a timescale of tens of minutes^19–21^. Making use of the simultaneous two-color imaging capability of MLS-SIM, we imaged both neural morphologies and PSD-95 distributions in awake mice brains.

We co-expressed membrane targeting mRuby3 and PSD-95 targeting fibronectin intrabody PSD95.FingR-EGFP^45^ in sparse populations of neurons in mice brains by virus injections. As in previous studies by EM and STED, MLS-SIM results reveal similar morphological diversity of PSD-95, ranging from perforated lines to complex patterns^19^ (Figure 5A and Video S7). They distributed differently on dendrite trunks and inside dendritic spines (Figure 5B): small PSD-95 structures were more commonly found on dendrite trunks, while elongated ones or those with complicated multi-domains were located predominately inside spine heads (Figure 5C). On the dendrite trunk, one prominent feature of PSD-95’s distribution was their tendency to be near the roots of filopodia (Figure 5D), which agreed with previous in vitro imaging studies^46^. MLS-SIM results further revealed small protrusions about 250 nm in height on average (Figure 5E), which could be very challenging for conventional microscopes to see, emerging within 500 nm from many PSD-95 domains on the dendrite trunk. These observations might underrepresent the overall coincidence of PSD-95 and small protrusions over longer time scales because we could have easily missed many other such events based on the highly dynamic nature of the growing and retracting of small protrusions near the PSD-95 domains (Figure 5F). Furthermore, considering the difficulty in capturing such instances along the z direction, the coincidence of PSD-95 and small protrusions might be even more prevalent, thus suggesting possible spatial correlation between them and connections between cellular preparation for filopodia emergence and PSD-95 recruitment.

**Figure 5.**
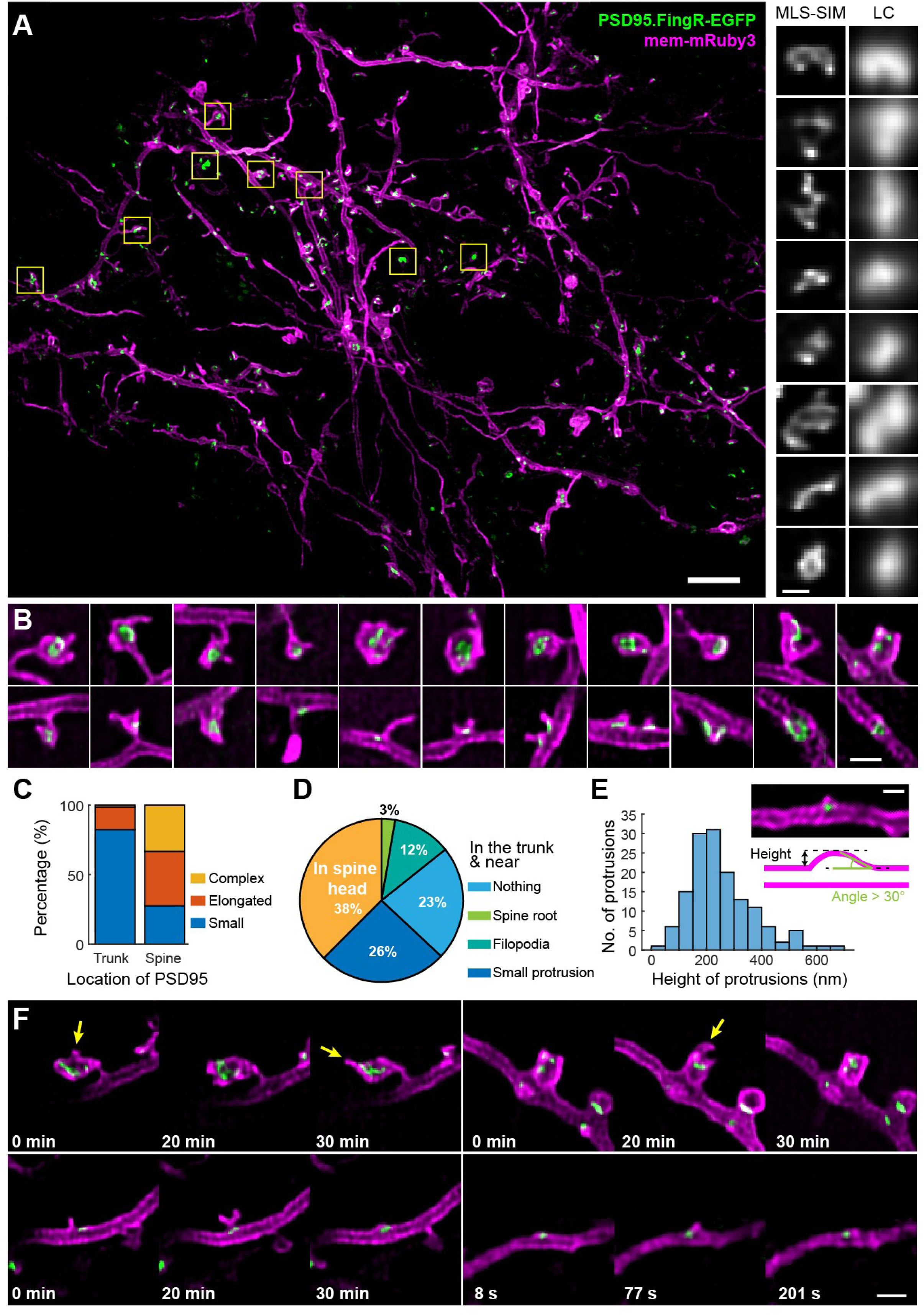
Two-color imaging of PSD-95 nanodomains and dendrite morphologies with MLS-SIM in awake mice brains. (A) Overview (left) of excitatory neurons expressing membrane-targeted mRuby3 and PSD-95 nanodomains labeled by EGFP. Magnified views (right) of PSD-95 in the regions indicated by the boxes imaged by MLS-SIM and conventional line confocal (LC) microscopy. (B) Examples of PSD-95 with diverse morphologies and localizations on the dendrites. (C) Statistics of differently shaped PSD-95 clusters, one for those on the dendrite trunks and the other for those inside the dendritic spines. (D) Statistics of different types of emerging locations of PSD-95 in all identified PSD-95 instances (n=560). (E) Histogram of the heights of small protrusions located on the dendrite trunk and alongside PSD-95 clusters. Inset: an example of a small protrusion and a schematic for how the protrusions were identified and their heights were measured. (F) Examples of time-lapse imaging of PSD-95 and dendrite. Dynamic growing and retracting of spinules (yellow arrow) on the spine and filopodia on the dendrite trunk were successfully captured. Scale bars: 5μm in (A, left panel); 200 nm in (A, right panel); 1μm in (B) and (F); 500 nm in (E, inset).

### Increasing imaging resolution by nonlinear MLS-SIM

Similar to nonlinear SIM^24,25^, the imaging resolution can be further improved in MLS-SIM by using nonlinear effects in fluorescence excitations. We chose saturated fluorescence excitation due to its simplicity and compatibility with MLS-SIM. Under saturated pattern illumination, the width of the minima, as opposed to the maxima, decreases with higher saturation effect^24,25^. Taking this into account, we redesigned the illumination patterns specifically for nonlinear MLS-SIM. In the first illumination line pattern responsible for to enhancing the y resolution, an intensity trough was introduced in the center of the line (Figure 6A). In the second and periodic pattern, two-beam replaced three-beam illumination to improve the x resolution (Figure 6A). Using a pulsed laser to concentrate the excitation energy within the fluorescence life time, saturation effect was achieved and resulted in sharper valleys around intensity minima in the excitation patterns and afforded extra resolution enhancement compared to those under linear excitation (Figure 6A). The resolution enhancement depends on the level of saturation. A higher level of saturation could in principle lead to higher resolution enhancement, but it might also reduce signal-to-background ratio (SBR) when imaging into thick tissues because the out-of-focus fluorescence was not generated under saturated excitations and increased linearly and thus faster than in-focus signal as saturation level increased (Figure 6B). In practice, we can finetune the saturation level for each sample based on the SBR and calibrated relationship between laser pulse energies and fluorescence emission rates (Figure 6B). To demonstrate the practical resolution enhancement by nonlinear MLS-SIM, we again imaged neurons expressing membrane-targeted EYFP in awake mouse brain. Thin neurites can be resolved as hollow tubes as narrow as about 100 nm in x-y (Figure 6C). Since the fluorescence lifetime (∼ns) is more than 10,000 times shorter than the line exposure time (∼100 us)^25^, enough number of laser pulses could be integrated during the acquisition of one line scan to allow imaging at high speed without compromising signal-to-noise ratio. In three successive frames captured on an awake mouse, sample motions led to marked image shifts and distortions (Figure 6D). However, high-resolution features could still be resolved in all frames (Figure 6D). The enhanced resolution due to the saturation effect inevitably came at the price of more severe photobleaching. Therefore, nonlinear MLS-SIM demonstrated here can be a better fit for applications where highest possible spatial resolution is more essential than longitudinal imaging capability.

**Figure 6.**
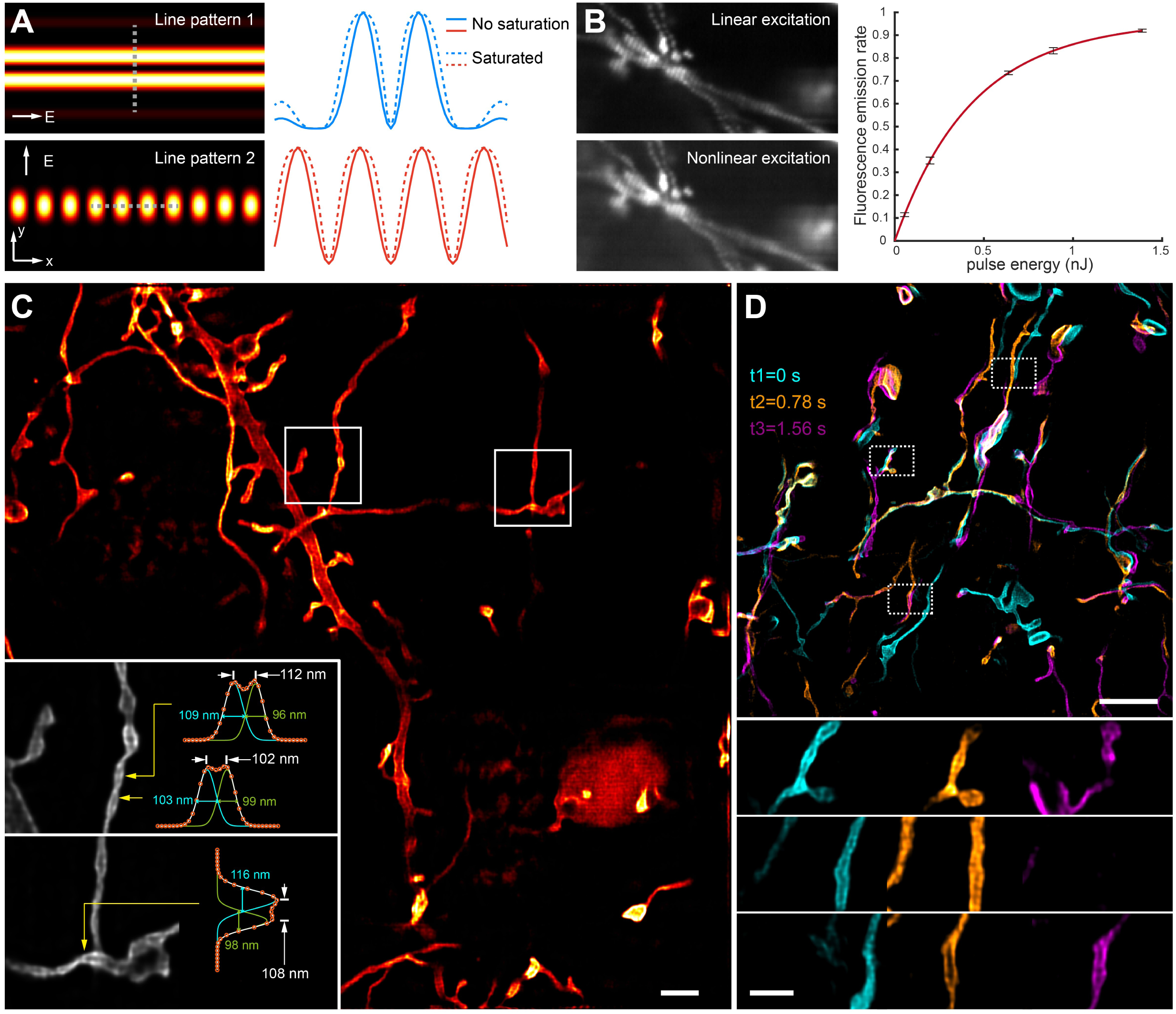
Nonlinear MLS-SIM imaging in awake mice brains. (A) Line excitation patterns used in nonlinear MLS-SIM and their intensity line profiles at indicated locations. Under saturated fluorescence excitation, effective fluorescence excitation patterns form steeper valleys around zero intensity nodes. (B) Comparison of SBR under pulsed laser excitations with different pulse energies. Based on a model describing saturation effect, the fluorescence emission rate at different laser pulse energies can be estimated experimentally. Error bars represent standard deviations of the fitted fluorescence emission rate measured on 500 selected imaging pixels in a calibrating imaging experiment (STAR Methods). (C) Nonlinear MLS-SIM imaging results of neurons expressing membrane-targeted EYFP in awake mouse brain. Insets: magnified views of features indicated by the boxes. Intensity line profiles are fitted by Gaussian functions to estimate the sizes and separations of the resolvable features, indicating the achievable practical resolution by nonlinear MLS-SIM. (D) Time-lapse imaging by nonlinear MLS-SIM in awake mouse brain. Three time points are colored differently. Due to the sample movement, images were distorted during the imaging, but three representative features (bottom) can still be resolved at super-resolution in these overall distorted images. Scale bars: 2 μm in (C); 5 μm in (D, top panel); 1μm in (D, bottom panels).

## DISCUSSION

It has been very challenging to image in super-resolution awake and behaving animals, especially in the presence of strong sample movement. MLS-SIM addressed this challenge and demonstrated its compatibility with biology, mainly benefitting from its mild photobleaching and phototoxicity, by volumetric and longitudinal imaging of the same structure over hundreds of time points and sometimes over multiple days. One of many promising applications is to image the brain because neural dynamics are profoundly affected by animals’ behavioral states. From imaging results of MLS-SIM, we could appreciate great diversities in morphologies and dynamics of dendritic spines and axonal boutons in awake mice brains. We could also identify tiny neuronal structures such as spinules that were originally observed under EM. We were able to track their dynamics during sleep-wake cycles or chronically over multiple days. In addition to observational studies on neural morphologies, the dynamics of certain relevant molecules, such as PSD-95, could be investigated simultaneously by two-color super-resolution imaging. Moreover, the resolution of MLS-SIM was further enhanced by using the nonlinear saturation effect.

While MLS-SIM makes it a reality to image awake animals in super-resolution by rejecting background fluorescence and tolerating animal’s movement, its achievable resolution is slightly inferior to wide-field SIM. MLS-SIM extends the resolution gain in two lateral directions by multiplexing two illumination patterns. In comparison, wide-field SIM enhances resolution in three lateral directions and nonlinear SIM collects high-resolution information in even more directions^25^. In practice, this problem was partially compensated by the nonlinear reconstruction that enforced non-negativity constraint in MLS-SIM. In the future, multiplexing more illumination patterns and introducing more priors learned from samples in image reconstructions^47,48^ can better address this problem. In nonlinear MLS-SIM reported here, we only explored the saturation effect for its simplicity, but the high peak illumination intensity led to accelerated photobleaching. Therefore, it would be interesting to explore the compatibility with MLS-SIM of other nonlinear effects, such as photoactivation^49–51^ and stimulated-emission-depletion^52^, to enhance longitudinal imaging capability.

Another key innovation of MLS-SIM is its reconstruction algorithm, an indispensable component of the technique. It introduces a new framework in which multiple images with different effective PSFs can be synthesized from the raw structured-illumination images and then coherently fused into a complete super-resolution image even though each of them has only partial information. This algorithm is applicable to implement constrained image reconstruction in wide-field SIM, and can serve as a general reconstruction framework for different types of structured illumination schemes.

Further improvements on MLS-SIM can benefit from advances in sample labeling^53,54^, hardware implementation, and algorithm development^47,48,55,56^. MLS-SIM can also be applied beyond neural imaging to image other cellular dynamics in the brain^57^, and other organs or tissues, such as beating hearts and breathing lungs, where sample motions have been prohibiting successful super-resolution imaging.

## ACKNOWLEDGMENTS

We thank L. Shao for offering insightful critiques of our manuscript. We thank J.W. Zhou, H.T. Xu, Z.Y. Liu, D.Q. Liu, J. He and Y. Mu for valuable discussions on neural imaging and data analysis. We thank J. Cao for mice care. We thank L. Cong, P. Yu, Z.Q. Shi for suggestions on data analysis. We thank H.T. Cao and Y.J. Xiu for suggestions on data analysis on sleep states. We thank T.L. Chen for initial experiments on zebrafish. Y.J. Zhang, L. Bai, Y.C. Zhao, T.L. Zhang, L.C. Ye and K. Wang were supported by STI2030-Major Projects (2021ZD0204503), National Natural Science Foundation of China (32125020). X. Wang and X.F. Du were supported by National Key R&D Program of China (2018YFA0801000 and 2018YFA0801001).

## AUTHOR CONTRIBUTIONS

K.W. conceptualized the project. Y.J.Z. built the MLS-SIM and wrote the code for image reconstructions and simulations. Y.J.Z. and X.W. performed zebrafish-related experiments under the supervision of X.F.D. and J.L.D.. Y.J.Z. performed mice-related experiments. Y.J.Z. performed EEG and EMG recording and data analysis on sleep states of mice under the supervision of Z.Z.. Y.J.Z., L.B., Y.C.Z., T.L.Z. and L.C.Y. contributed to data analysis and discussion. K.W. and Y.J.Z. wrote the manuscript with inputs from all authors. K.W. and J.L.D. supervised the project.

## SUPPLEMENTARY FIGURE TITLES AND LEGENDS

**Figure S1. Experimental implementation and numerical simulations of MLS-SIM, related to** **Figure 1** **and STAR Methods**

A. Schematics of the optical system of MLS-SIM. DM, dichroic mirror; PBS, polarizing beam splitter; AOTF, acousto-optic tunable filter; RPG, rotation phase grating; PD, photodetector; TF, transmission filter; SM, scanning mirror; PM, phase mask; SF, spatial filter. PM and SF are absent in linear MLS-SIM imaging and present only in nonlinear MLS-SIM imaging.
B. Schematics of device synchronization. During the capture of a single frame, the camera is exposed multiple times and each exposure corresponds to one line in the resulting imaging frame. Two acousto-optic tunable filters (AOTF1 and AOTF2 in (A)) switch two different patterns on and off alternatively. In two-color imaging, two different colors are interleaved before interleaving patterns to ensure co-alignment between the two colors. Y scanning mirror (SM in (A)) performs a single continuous scan in the y direction in each frame. At the same time, the phase grating (RPG in (A)) is continuously rotated to scan the second excitation pattern along the x direction. Due to the instability of the employed rotation stage, the rotational speed of the phase grating is monitored by a photodetector (PD in (A)) for phase estimation of the second illumination pattern in image reconstruction.
C. Phase modulation mask for structured line illumination generation. Left, Schematics of the circularly distributed grating (RPG in (A)). The blue line on the left part of the grating indicates the location where the line excitation laser is focused on. The green line on the magnified view of the grating indicates the location of the cross-section view shown at the bottom. It is a stepped transmission phase grating fabricated on the fused silica wafer. Right, A photo of the customized phase grating. The bottom right is a magnified photo taken on a differential interference contrast (DIC) microscope.
D. Experimental implementation of MLS-SIM. Top, overview of the optical system. The yellow line indicates the light path of the excitation beam. Bottom, a magnified view of the multiplexing of two excitation patterns. The red line indicates the light path of the 635 nm laser for rotation phase grating monitoring. All labels of optical elements are the same as their counterparts described in (A).
E. Numerical simulations of line excitation patterns employed in linear and nonlinear MLS-SIM and their comparison with conventional diffraction-limited focus spot. First row, schematics of different patterns of excitations on the imaging objective’s pupil plane. Second row, 3D views of simulated intensity distributions of different excitation patterns near the focal plane. Third row, comparisons of intensity profiles at indicated locations in the excitation patterns. In the simulation, the wavelength of the excitation laser is 488 nm, and the numerical aperture of the objective is 1.15. Polarizations are indicated by “E” and full vector model was employed in the simulation^58^. Among all three patterns, line patterns 1 and 2 have the tightest focuses in the y and x directions, respectively. Line pattern 2 has the tightest focus in the z direction. The effective excitation profiles of y4 and x5 in nonlinear excitation conditions assume a fluorescence emission rate^25^ of 0.9.

**Figure S2. Schematics of image acquisition and reconstruction in MLS-SIM, related to STAR Methods**

A. Simplified schematics of consecutive 10 camera frames (Frame1–Frame10) captured under alternative excitations of two different patterns during scanning in y direction at 10 different locations (y1-y10). Each camera frame has two rows and each row has 10 imaging pixels. To avoid confusion, imaging pixels are termed as detectors hereafter. The detector measurement M(i,j,k) can be indexed by the column number (i), row number (j) and frame number (k). Depending on the spatial relationships between excitation patterns and detectors, the detector measurements are divided into eight different groups. Each detector measurement belongs to a certain group indicated by the index on the pixel in the schematics.
B. Schematics of super-resolution image reconstruction. Each group of detector measurements is captured under the same excitation pattern from the perspectives of these detectors. They share the same effective PSF (PSFEF), which depends on the spatial relationship between excitation PSF (PSFExt) and detection PSF (PSFDet). These detector measurements can be stitched into 2D images (CFImg) consisting of 10 rows by 10 columns of pixels. Not all pixels in the 2D array are provided as excitation patterns are interleaved in time and space. These pre-processed images CFImg are up-sampled first along the x direction by filling in unsampled pixels. The up-sampled images USCFImg are then jointly deconvolved using a customized algorithm derived from Richardson-Lucy deconvolution. Finally, pixel binning along the y axis is performed on the deconvolved super-resolution image to make the final image have the same scales along the x and y directions.

**Figure. S3. Experimental characterizations on motion tolerance and long-term volumetric imaging capabilities of MLS-SIM, related to Figure 2**

A. Resolution characterization of MLS-SIM on static fluorescent beads with diameters of 100 nm.
B. Reconstructions of the same group of representative fluorescent beads moving at three different speeds (25, 50, and 100 micrometers per second) and in eight different directions.
C. Statistical quantifications of experimentally obtained resolutions by resolving bead pairs with different orientations.
D. MIP of an imaging stack captured in an awake mouse brain.
E. Magnified view of the indicated ROI in (D) and the intensity line profiles across the time series at the location indicated by “X”. Small features could be resolved during the entire time series.
F. Change of total fluorescence collected in the image stacks across time. An abrupt increase in total fluorescence was caused by the intentional increase of excitation laser power to maintain imaging quality.

**Figure S4. Numerical simulations and experimental characterizations on multi-layer-reconstruction in MLS-SIM, related to STAR Methods**

A. Diversities of detector groups and their corresponding effective PSFs employed in simulations. Ten images were collected with 10 different PSFs (Principles of imaging and reconstruction in MLS-SIM, STAR Methods). Top, 10 different spatial relationships between excitation patterns and detectors on the camera generate 10 different effective PSFs. Middle, effective PSFs at different depths from the focal plane. Bottom, the corresponding optical transfer functions (OTFs) of these effective PSFs.
B. Simulations on noise amplification in multi-layer reconstruction. Based on a multi-layer reconstruction model in spatial frequency space, the reliability of reconstruction could be estimated by the noise amplification factors. The noise amplification factors in the spatial frequency space of two reconstructed layers in two configurations are presented on the left and right, respectively. Dashed circles indicate spatial frequencies supporting indicated lateral resolutions.
C. Representative experimental super-resolution reconstruction using single-layer and two different multi-layer reconstruction strategies. Multi-layer reconstructions yielded images with reduced out-of-focus backgrounds. The performance of multi-layer reconstructions depended on the choices of reconstructed layers. Three-layer reconstruction at z=0 and z=±150nm (middle) produce more reconstruction artifacts than the reconstruction at layers z=0 and z=±450nm (right).

## STAR METHODS

### RESOURCE AVAILABILITY

#### Lead contact

Further information and requests for resources and reagents should be directed to and will be fulfilled by the Lead Contact, Kai Wang.

#### Materials availability

All reagents generated in this study are available from the lead contact.

#### Data and code availability

- All imaging raw data generated in this study will be shared by the lead contact upon request.
- All original code and example data has been deposited at Science Data Bank and is publicly available as of the date of publication. DOIs are listed in the key resources table.
- Any additional information required to reanalyze the data reported in this paper is available from the lead contact upon request.

### EXPERIMENTAL MODEL AND SUBJECT DETAILS

#### Zebrafish

Experiments involving zebrafish were performed in accordance with the institutional guidelines approved by the Animal Care and Use Committee of the Center for Excellence in Brain Science and Intelligence Technology, Chinese Academy of Sciences (NA-046-2023). Adult zebrafish were maintained in the National Zebrafish Resources of China (Shanghai, China) with an automatic fish housing system at 28°C. Zebrafish embryos and larvae were raised in 10% Hank’s solution (in mM, 140 NaCl, 5.4 KCl, 0.25 Na2HPO4, 0.44 KH2PO4, 1.3 CaCl2, 1.0 MgSO4, and 4.2 NaHCO3, pH 7.2) at 28°C. The less pigmented *nacre* mutants (*mitfa^w^*^2^)^59^ with transient transgene expression at 4 - 5 days post-fertilization (dpf) was used for the study. The sex of the larvae at this stage is not determined.

#### Mice

All experimental procedures involving the usage of mice were approved by the Animal Care and Use Committee of the Center for Excellence in Brain Science and Intelligence Technology, Chinese Academy of Sciences. For morphological imaging of dendrites, 10-week-old male C57BL/6J mice were used. For two-color imaging experiments, 6-week-old male C57BL/6J mice were used. For morphological imaging of axonal boutons of interneurons, adult male or female Sst-Cre mice were used. For nonlinear MLS-SIM imaging experiments, adult male or female Sst-Cre mice were used. For imaging during sleep, mice were housed under standardized conditions with a 12 h–12 h light–dark cycle (light phase: 07:00 to 19:00), with the temperature controlled at 22–23 °C and humidity at 40–70%. For all other experiments, mice were housed under the same condition but with an inverted light-dark cycle (light phase: 22:00 to 10:00). All imaging experiments were carried out in daytime. The expression of fluorescent proteins was mediated by stereotaxic adeno-associated virus (AAV) injection. For morphological imaging of dendrites, AAV2/9-CamKII-Cre and AAV2/9-hSyn-DIO-mem-EGFP were injected in C57BL/6J mice. For morphological imaging of axonal boutons of interneurons, AAV2/9-hSyn-DIO-mem-EGFP was injected in Sst-Cre mice. For nonlinear MLS-SIM imaging experiments, AAV2/9-hSyn-DIO-mem-EYFP was injected in Sst-Cre mice. For two-color imaging of PSD-95 and neuron morphologies, AAV2/9-CamKII-Cre, AAV2/9-hSyn-DIO-mem-mRuby3, and AAV2/8-EF1A-DIO-PSD95.FingR-EGFP-CCR5TC^60^ were co-injected in C57BL/6J mice.

### METHOD DETAILS

#### Experimental setup of MLS-SIM

The imaging system (Figures S1A to S1D) was based on a platform of confocal line scanning microscope. In the laser excitation light path, two laser beams (488 nm and 561 nm, OBIS LS 488-100 and OBIS LS 561-150, Coherent) with their polarizations independently controlled by two half-wave plates WP1 and WP2 (GCL-060811, Daheng Optics) were combined by a dichroic mirror DM1 (Di02-R488, Semrock). The combined laser beam was then split into two beams with orthogonal polarizations by polarizing beam split PBS1 (GCC-402103, Daheng Optics). Two acoustic optical tunable filters AOTF1 and AOTF2 (97-03151-01, G&H) controlled the transmission of wavelength and power of these laser beams at high speed. After AOTF1, the y-polarized laser beam was first expanded and collimated to 10 mm in diameter by a pair of lenses L1 & L2 (f1=10 mm & f2=100 mm, AC060-010-A and AC254-100-A, Thorlabs) and then focused into a line along the x direction by the cylindrical lens L3 (f3=150 mm, ACY254-150-A, Thorlabs). A transmission phase grating RPG (Figure S1C) mounted on a rotation stage (DDR25/M, Thorlabs) was placed at the focal plane of this line focus. The phase grating was a binary stepped grating fabricated on a 100 mm diameter fused silica wafer with 6,600 circularly distributed periods. The depth of the grating was 300 nm to provide 138 nm optical path difference. We focused the laser line perpendicular to the grating line and 38 mm from the center of the grating plate (Figure S1C). In this case, the phase of the focused laser line was modulated with a period of ∼36 µm. This period could be fine-tuned by adjusting the distance between the line focus and the center of the grating plate. Since the rotational speed of the phase grating could not be controlled with high precision, we tracked the rotation of the grating by focusing a 635 nm laser beam (3 mm diameter, CPS635R, Thorlabs) at the center of the line focus of the excitation laser on the phase grating. The phase modulation of the focused 635 nm laser beam changed the intensity distribution of the transmitted beam. The transmitted beam was spatially filtered by an iris and monitored by a phase grating monitoring photodetector (PD). When phase grating rotates, the periodic modulation of the 635 nm laser beam could be detected by PD (Figure S1B). The phase-modulated line focus of the excitation laser beam is then relayed to the sample plane by three pairs of lenses that were all arranged in 4-f configurations: including L5 & L6 (f5=100 mm and f6=100 mm, AC254-100-A, Thorlabs), L7 & L8 (f7=275 mm and f8=175 mm, 47-644 and 47-648, Edmund Optics) and L9 & Objective (f9= 200 mm, AC254-200-A, Thorlabs, and NA=1.15, 40×, CFI Apo LWD Lambda S 40XC WI, Nikon). In the other light path after PBS1 and AOTF2, the x-polarized laser beam was also expanded by L10 & L11 (f10=10 mm & f11=100 mm, AC060-010-A and AC254-100-A, Thorlabs) and then focused into a line along the x direction by a cylindrical lens L12 (f12=150 mm, ACY254-150-A, Thorlabs). This x-polarized beam was recombined with y-polarized beam by polarizing beam split PBS2 (GCC-402103, Daheng Optics) after passing through a relay lens L13 (f13=100 mm, AC254-100-A, Thorlabs), which was also configured as a 4-f system together with L6. During the relay of the excitation laser beams to the imaging objective, a two-axis piezo scanning mirror SM (PT2M25-2000S-S, Nano Motions Technology Co. Ltd.) was placed between L7 and L8 and conjugated to the back pupil of the imaging objective for beam scanning. Fluorescence detection shared the same light path between the imaging objective and the relay lens L7. The fluorescence signal collected by the imaging objective was de-scanned by SM and focused on to the imaging camera (C15440-20UP, Hamamatsu) by L7. The excitation and detection light paths were separated by a dichroic mirror DM2 (Di03-R488/561, Semrock). A multiband transmission filter TF (Em01-R488/568, Semrock) was inserted in front of the camera to block excitation laser beams. A slim region of interest (ROI) consisting 4 or 8 lines of pixels near the center of the imaging sensor on camera were selected to collect images at high speed (up to 20,000 Hz).

The nonlinear version of the system was slightly modified on its linear version (Figure S1A). First, the excitation laser was changed to a pulsed laser (FL-515-Pico 1W, Changchun New Industries Optoelectronics Tech. Co., Ltd.). It generated 600 ps pulses with a repetition rate of 1 MHz and at the wavelength of 515 nm. Second, DM2 (ZT442/514/561rpc, Chroma) and TF (ZET442/514/561m-TRF, Chroma) were changed to match the requirement for fluorescence excitation and detection. Third, the phase grating RPG in y-polarized excitation light path had a different phase modulation depth (288 nm optical path difference) and a spatial filtering mask was placed at the pupil plane of L6 to block zero order diffraction beam and generate a two-beam interfering excitation pattern (Figure S1A). Fourth, an additional phase mask PM (Figure S1A) generating hollow line excitation pattern (Figure S1E) was placed between L11 and L12.

During the imaging acquisition, the rotational speed of the phase grating was set to match the imaging speed. We empirically set the rotational speed of the phase grating to introduce 2π phase shift during the acquisition of every 12 lines, comprising 6 lines under first and second excitation patterns. In nonlinear MLS-SIM imaging, a 2π phase shift of the second excitation pattern was achieved during the acquisition of every 14 lines. The piezo scanning mirror SM, imaging camera, photodetector PD, AOTF1 and AOTF2 were properly synchronized during one-color and two-color imaging experiments (Figure S1B).

### Principles of imaging and reconstruction in MLS-SIM

#### (1)#Forward imaging model of MLS-SIM

The forward imaging model describes how MLS-SIM performs imaging and collects raw data from samples. In particular, the collected raw data can be properly re-arranged into multiple confocal images that can be jointly deconvolved later to reconstruct super-resolution images.

In MLS-SIM, imaging pixels on the camera works as an arrayed detector and each detector can be indexed by its location on the camera 𝐷(𝑥, 𝑦). The detection point spread function and the patterned excitation beam profile can also be described in this coordinate on the camera as 𝑃𝑆𝐹_𝐷𝑒𝑡_(𝑥, 𝑦) and 𝑃𝑆𝐹_𝐸𝑥𝑡_(𝑥, 𝑦), respectively. When the excitation beam is scanned to location (𝑥_𝑠_, 𝑦_𝑠_) on the sample, the imaging camera is also shifted to this location and the sample can be written as 𝑂𝑏𝑗(𝑥 + 𝑥_𝑠_, 𝑦 + 𝑦_𝑠_) in the coordinate on the camera. At this location (𝑥_𝑠_, 𝑦_𝑠_), the camera captures an image, which can be written as:

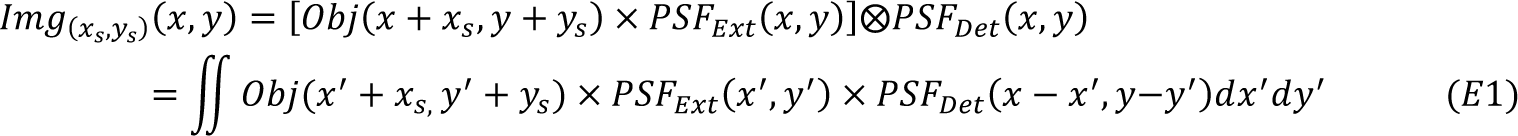

where ⊗ represents convolution operation.

Let’s assume we can do 2D scanning (𝑥_𝑠_, 𝑦_𝑠_) over the sample. In this case, each detector 𝐷(𝑥 = 𝑥_𝑑_, 𝑦 = 𝑦_𝑑_) on the camera works as a confocal microscope and can produce a 2D image 𝐶𝐹𝐼𝑚𝑔_𝐷(𝑥𝑑,𝑦𝑑)_(𝑥_𝑠_, 𝑦_𝑠_) . According to equation E1, the confocal image 𝐶𝐹𝐼𝑚𝑔_𝐷(𝑥𝑑,𝑦𝑑)_(𝑥_𝑠_, 𝑦_𝑠_) can be written as:

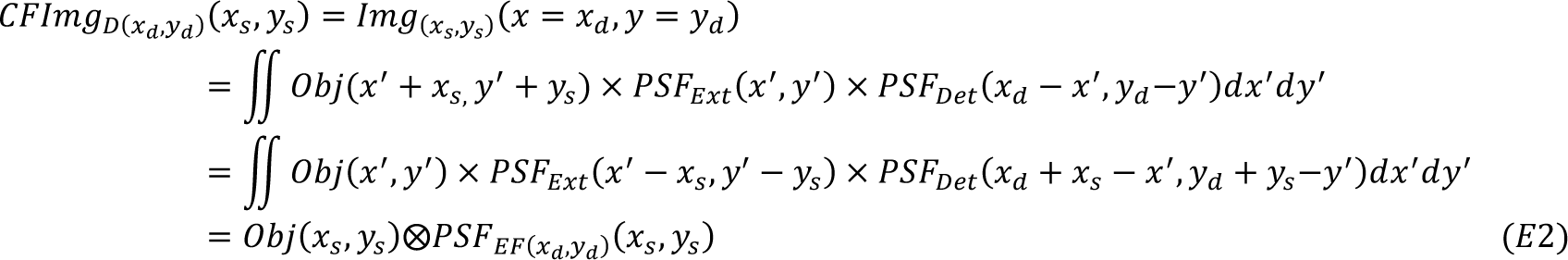

Where 𝑃𝑆𝐹_𝐸𝐹(𝑥𝑑,𝑦𝑑)_ is the effective point spread function and defined as:

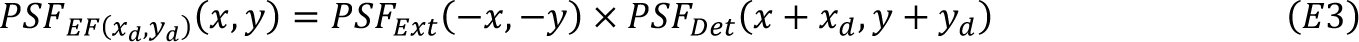

In MLS-SIM imaging, we only perform 1D scan along y axis. However, we will show that similar 2D confocal images 𝐶𝐹𝐼𝑚𝑔_𝐷(𝑥𝑑,𝑦𝑑)_(𝑥_𝑠_, 𝑦_𝑠_) can also be formed during this process by making use of the translational invariance of the first illumination pattern and periodicity of the second illumination pattern.

To simplify the discussion of the model, we assume that camera only has an array of 10×2 detectors (Figure S2A) and makes totally 10 × 2 × 10 measurements 𝑀(𝑖, 𝑗, 𝑘) (1 ≤ 𝑖 ≤ 10,1 ≤ 𝑗 ≤ 2,1 ≤ 𝑘 ≤ 10,) during a scan along y direction at 10 different locations (𝑦_1_, 𝑦_2_ … 𝑦_10_). Two illumination patterns are interleaved one after the other during the scan. Since the effective PSF only depends on the spatial relationship between the illumination pattern and the pixel detector (equation E3), we can see that many measurements from different detectors and in different frames share same effective PSFs. In fact, we can divide these measurements into 8 different groups (Figure S2B). Under the first illumination pattern, there are only two different effective PSFs (Figure S2B), because the excitation patterns are the same from the point of views of all detectors in the same row. In this case, the measurements made by detectors in the same row but different columns in a frame form a row of pixels in the 2D image that is traditionally formed during a 2D scan as discussed above. Therefore, 1D scan in y direction of this row of pixels yields a 2D confocal image (Figure S2B). Because only 5 frames are captured under first illumination pattern, the corresponding 5 out of 10 rows in the 2D confocal image are filled with measurements. The second illumination pattern is more complex than the first one because its intensity varies along both x and y directions. To account for all different situations, we need to define 6 different effective PSFs (Figure S2B). In addition, the phase of the periodic pattern along x axis is changing from frame to frame. Therefore, the same pixel detector on the camera makes measurements under different effective PSFs in different frames. However, we can correctly assign these measurements into different confocal images if we know the phase of the excitation pattern in each frame. In the end, we generate 8 confocal images 𝐶𝐹𝐼𝑚𝑔_𝐷𝑘_(𝑥_𝑠_, 𝑦_𝑠_) (1 ≤ 𝑘 ≤ 8) whose pixels are only partially sampled. Each confocal image can be considered as being collected through 2D scanning under its corresponding effective PSF 𝑃𝑆𝐹_𝐸𝐹𝑘_ (𝑥_𝑠_, 𝑦_𝑠_) (1 ≤ 𝑘 ≤ 8).

#### (2)#Super-resolution image reconstruction of MLS-SIM

The reconstruction algorithm combines partially sampled confocal images described above into one fully sampled super-resolution image. The whole process mainly consists of three steps (Figure S2B): (a) Image up-sampling, (b) Joint deconvolution and (c) Pixel Binning.

#### (2a)#Image up-sampling

To prepare for joint deconvolution, confocal images 𝐶𝐹𝐼𝑚𝑔_𝐷𝑘_(𝑥_𝑠_, 𝑦_𝑠_) are firstly up-sampled to make sure the pixel size of the deconvolved image is small enough to satisfy Nyquist sampling criterion for super-resolution imaging. The pixel size of 𝐶𝐹𝐼𝑚𝑔_𝐷𝑘_(𝑥_𝑠_, 𝑦_𝑠_) in y direction is determined by the scanning step size, which is usually set small to ensure that enough images of the same area are collected under the second illumination pattern with different phases. In many of our linear MLS-SIM imaging experiments, the scanning step size is 10 nm. The pixel size of 𝐶𝐹𝐼𝑚𝑔_𝐷𝑘_(𝑥_𝑠_, 𝑦_𝑠_) in x direction is the same as the camera’s pixel size (103 nm), which is designed to satisfy Nyquist sampling criterion for diffraction-limited imaging. Therefore, we only need to up-sample the image in x direction. To support resolution doubling in linear MLS-SIM, we half the pixel size in x direction. In nonlinear MLS-SIM, we further reduce the pixel size in both x and y directions to allow reconstruction at even higher resolution.

#### (2b)#Joint deconvolution

The up-sampled confocal images 𝑈𝑆𝐶𝐹𝐼𝑚𝑔_𝐷𝑘_(𝑥_𝑠_, 𝑦_𝑠_) contain mutually complementary information that can be jointly deconvolved by adapting Richardson-Lucy deconvolution algorithm to our framework. To prepare for the deconvolution, we define 𝑀𝑎𝑠𝑘_𝐷𝑘_(𝑥_𝑠_, 𝑦_𝑠_) for each of the 𝑈𝑆𝐶𝐹𝐼𝑚𝑔_𝐷𝑘_ (𝑥_𝑠_, 𝑦_𝑠_) to mark the sampled and unsampled pixels in 𝑈𝑆𝐶𝐹𝐼𝑚𝑔_𝐷𝑘_(𝑥_𝑠_, 𝑦_𝑠_) . 𝑀𝑎𝑠𝑘_𝐷𝑘_(𝑥_𝑠_, 𝑦_𝑠_) is a binary image and has the same size as 𝑈𝑆𝐶𝐹𝐼𝑚𝑔_𝐷𝑘_(𝑥_𝑠_, 𝑦_𝑠_). Pixels that are correspondingly sampled in 𝑈𝑆𝐶𝐹𝐼𝑚𝑔_𝐷𝑘_(𝑥_𝑠_, 𝑦_𝑠_) are set to the value of one in 𝑀𝑎𝑠𝑘_𝐷𝑘_(𝑥_𝑠_, 𝑦_𝑠_). All other pixels are set as zero. Unsampled pixels in 𝑈𝑆𝐶𝐹𝐼𝑚𝑔_𝐷𝑘_(𝑥_𝑠_, 𝑦_𝑠_) are also set as zero. The pseudo-code for joint deconvolution can be summarized below. Without causing confusion, notations (𝑥_𝑠_, 𝑦_𝑠_) is simplified as (𝑥, 𝑦) in the pseudo-code.

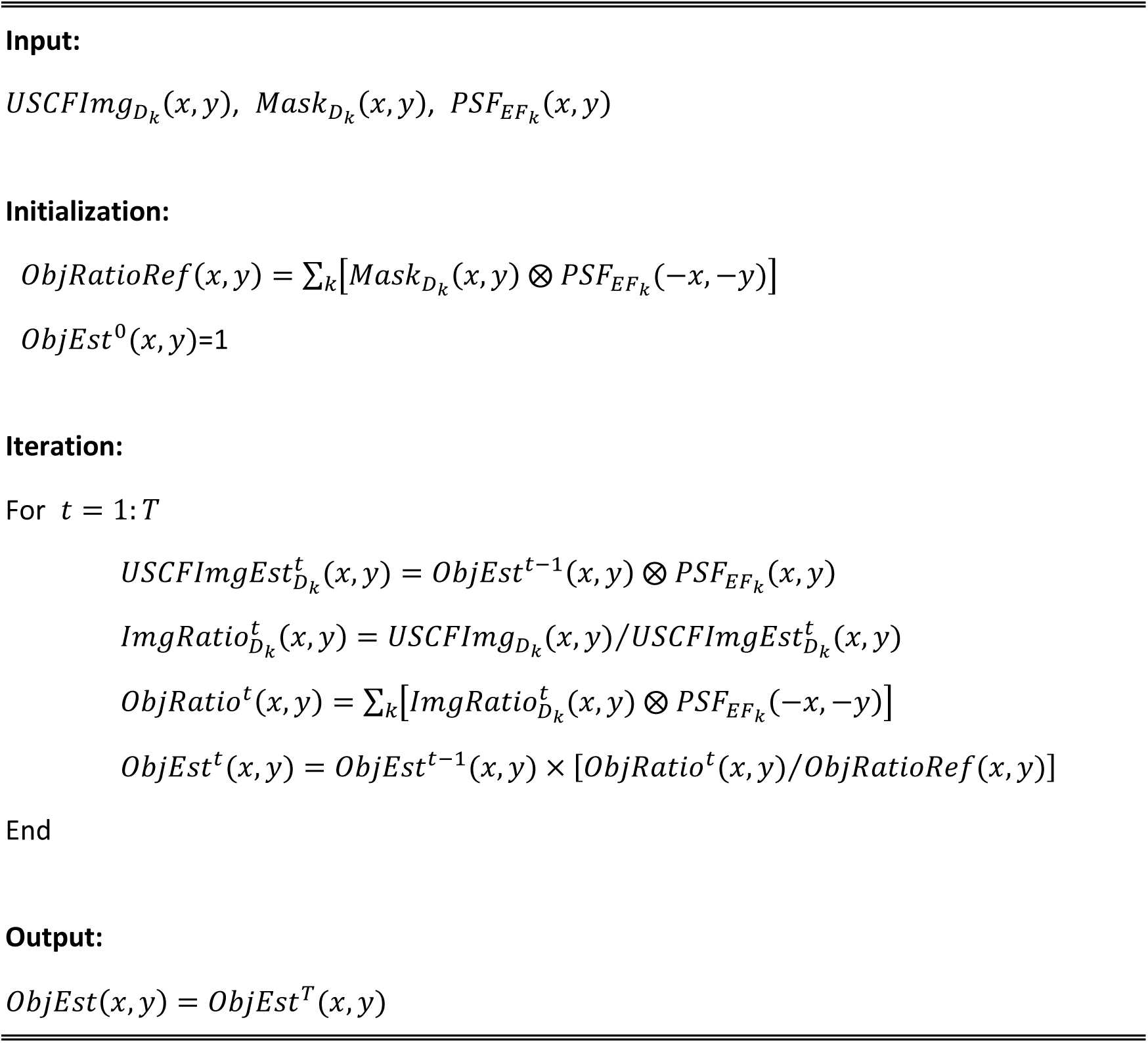

#### (2c)#Pixel binning

Joint deconvolution yields an estimation of the sample 𝑂𝑏𝑗𝐸𝑠𝑡, however, its pixel has different sizes in x and y directions. For example, in many of our linear MLS-SIM imaging experiments, the pixel size in y direction is only 10 nm. The algorithm above may render reconstructed images many details that are beyond the actual resolving capability of MLS-SIM. To avoid such confusion, we can bin neighbor pixels such that the resulting pixel size is just right to satisfy Nyquist sampling criterion for expected resolution. In practice, we bin 5 neighboring pixels in y direction to make resulting pixel has the same size (50 nm) in both x and y directions (Figure S2B).

#### (3)#3D model for imaging and reconstruction

Although MLS-SIM performs 2D imaging during a single scan, but the diversity of the excitation pattern may help discern information that is within or away from the focal plane. To take advantage of this fact, we need to build 3D model for imaging and reconstruction.

If we approximate the 3D sample by discrete layers of 2D structures 𝑂𝑏𝑗(𝑥, 𝑦, 𝑧) ≈ 𝑂𝑏𝑗(𝑥, 𝑦, 𝑧_𝑛_) (𝑛 = 0, ±1, …), where 𝑛 = 0 represents the focal plane. The forward imaging model (equation E2) can be updated as:

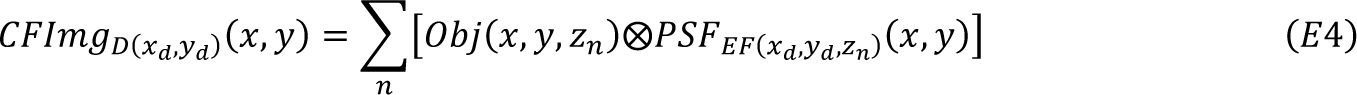

where 𝑃𝑆𝐹_𝐸𝐹(𝑥𝑑,𝑦𝑑,𝑧𝑛)_(𝑥, 𝑦) is defined as:

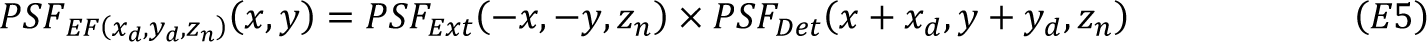

Based on this model, the corresponding joint deconvolution can be described as below. We adopt the same notions for different groups of PSFs in 2D and add subscript index 𝑛 to represent different layers.

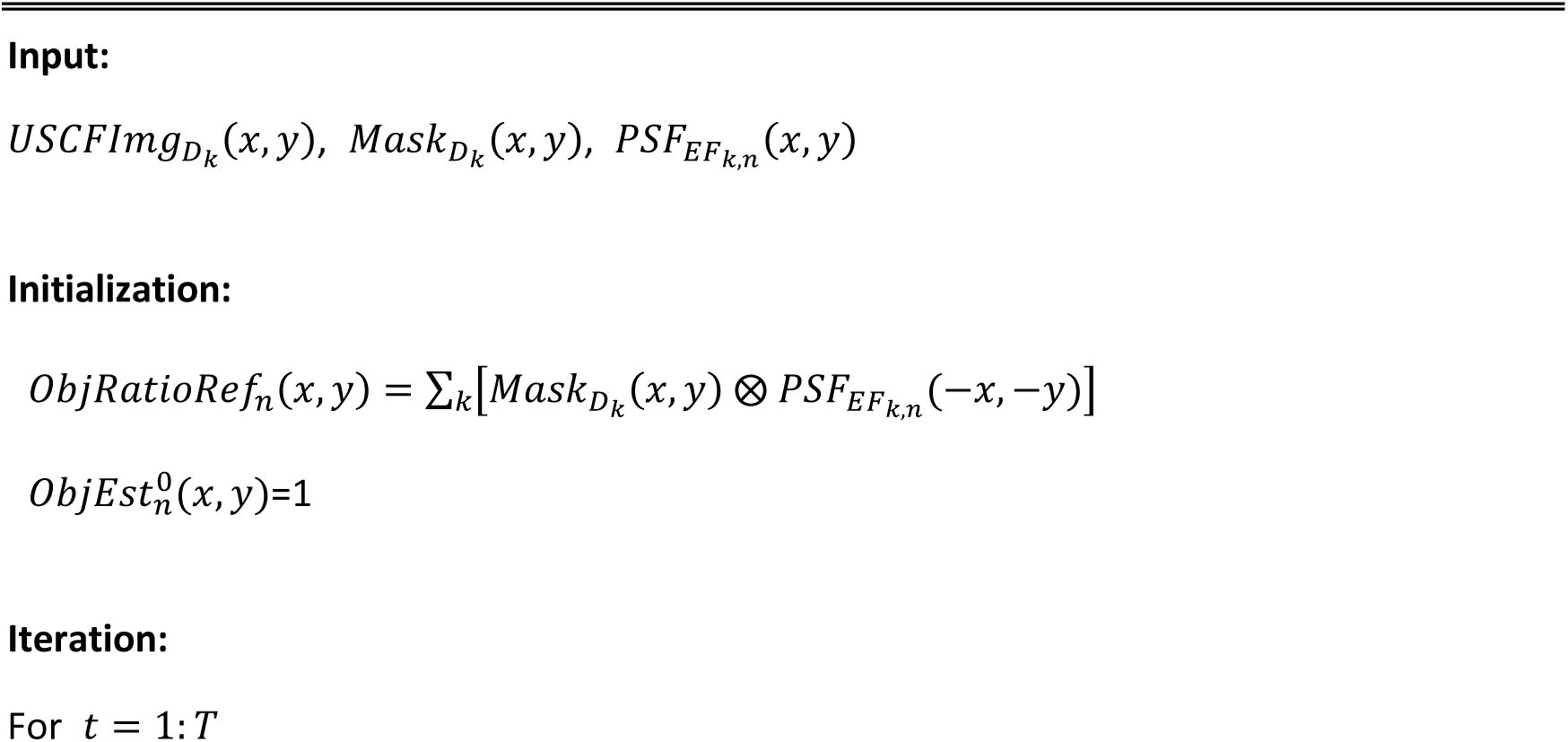
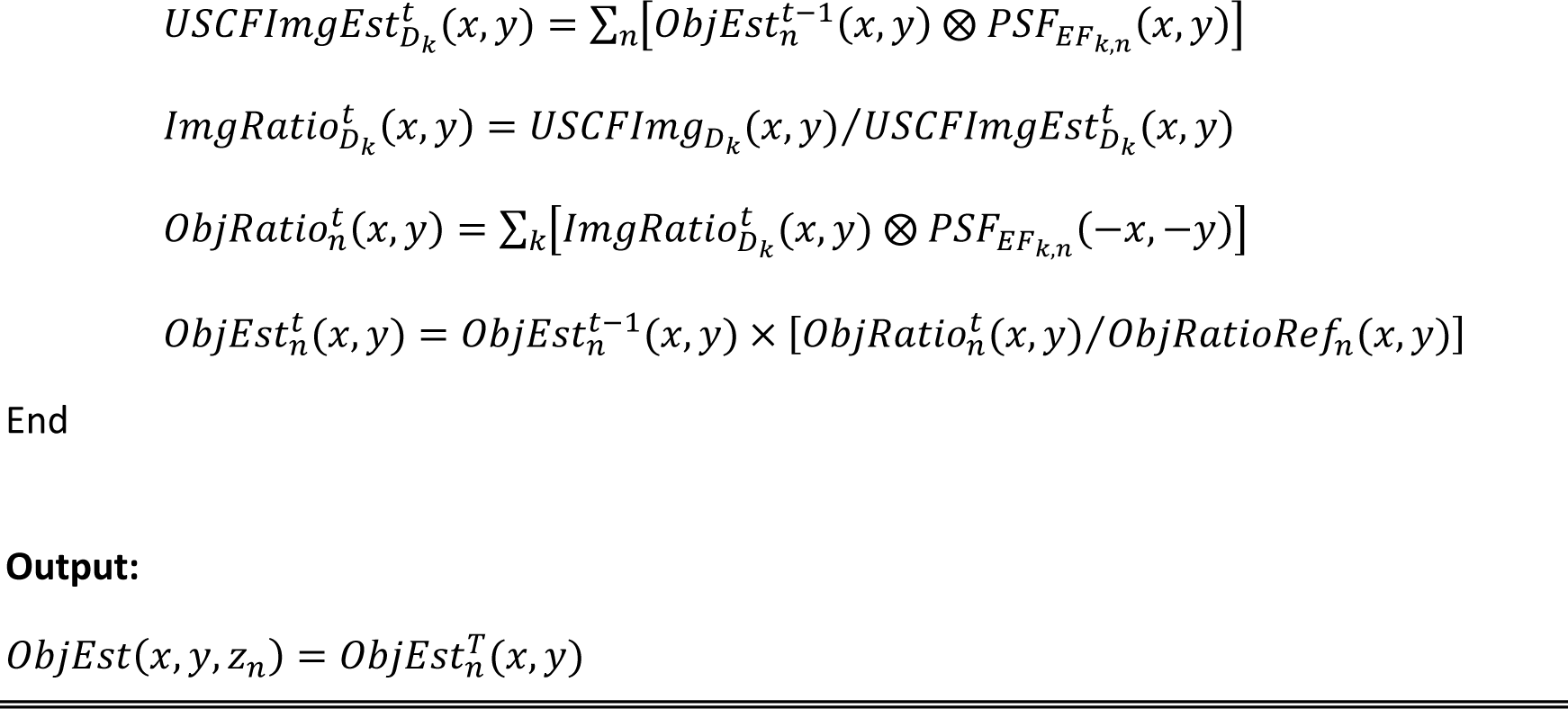

The above algorithm cannot guarantee a proper reconstruction if there are many layers. In practice, we only need to account for 3 layers (above, in and below the focal plane) in MLS-SIM because MLS-SIM employs line confocal detection scheme and can physically reject fluorescence that are far away from the focal plane. The goal of this joint deconvolution is not to reconstruct multiple layers simultaneously, but to reduce interference from out of focus fluorescence and achieve super-resolution optical sectioning for the in-focus layer.

Instead of presenting a formal proof, we provide an argument on the rationale for multi-layer reconstruction. To simplify the discussion, we assume all confocal images 𝐶𝐹𝐼𝑚𝑔_𝐷_𝑘_(𝑥, 𝑦) are fully sampled, which can be approximated in MLS-SIM by making y scanning step small and binning neighboring steps together. Since different confocal images 𝐶𝐹𝐼𝑚𝑔_𝐷_𝑘_(𝑥, 𝑦) are generated by different effective PSFs in the forward imaging model, the diversity of these effective PSFs may enable reconstruction of multi-layers at the same time. In summary, we want to reconstruct 𝑛 unknow layers 𝑂𝑏𝑗(𝑥, 𝑦, 𝑧_𝑛_) from 𝑘 measured images 𝐶𝐹𝐼𝑚𝑔_𝐷𝑘_(𝑥, 𝑦) and their relationships are described by the forward imaging model:

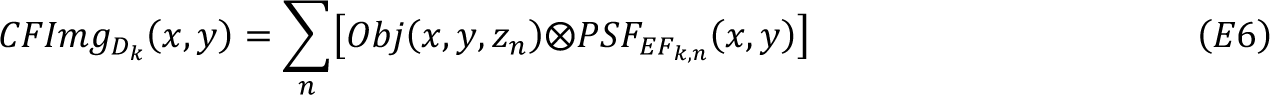

We can rewrite this set of equations in spatial frequency space as:

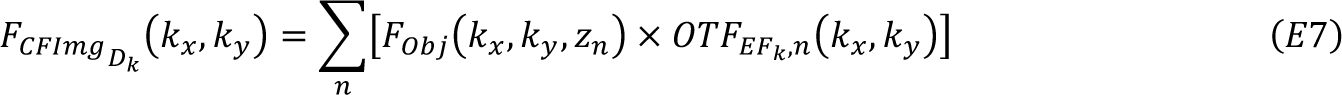

where𝐹_𝐶𝐹𝐼𝑚𝑔_ (𝑘_𝑥_, 𝑘_𝑦_), 𝐹_𝑂𝑏𝑗_(𝑘_𝑥_, 𝑘_𝑦_, 𝑧_𝑛_) and 𝑂𝑇𝐹_𝐸𝐹𝑘,𝑛_(𝑘_𝑥_, 𝑘_𝑦_) are Fourier transforms of 𝑘 𝐶𝐹𝐼𝑚𝑔_𝐷𝑘_(𝑥, 𝑦), 𝑂𝑏𝑗(𝑥, 𝑦, 𝑧_𝑛_) and 𝑃𝑆𝐹_𝐸𝐹𝑘,𝑛_ (𝑥, 𝑦), respectively.

For each spatial frequency component (𝑘_𝑥_, 𝑘_𝑦_), equation E7 can be written in matrix form as:

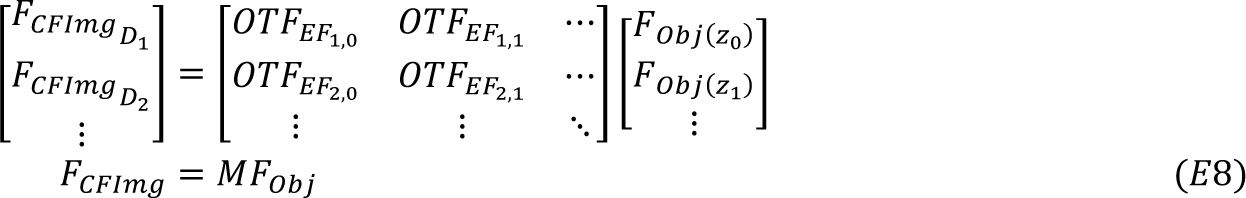

This set of equations is over-constrained if the number of measured confocal images (𝑘) is more than the number of layers (𝑛) we want to reconstruct. In this case, the least square estimation of the Fourier transforms of the object can be found by:

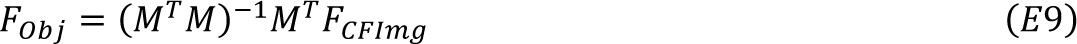

However, the equation E8 can be ill-conditioned depending on the properties of matrix 𝑀 and solutions found by equation E9 can be sensitive to noise. To test the reliability of solutions found by equation E9, we did a series of simulations to calculate the average noise amplification factor 𝐴𝐹_𝑁𝑜𝑖𝑠𝑒_ when feeding random 𝐹_𝐶𝐹𝐼𝑚𝑔_ with random noise 𝛿𝐹_𝐶𝐹𝐼𝑚𝑔_ to equation E9. The noise amplification factor 𝐴𝐹_𝑁𝑜𝑖𝑠𝑒_ is defined as:

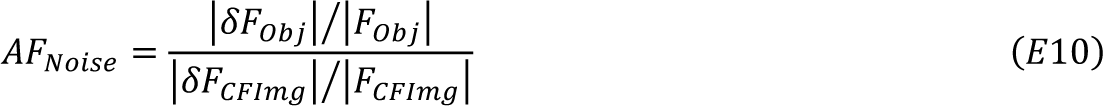

where (𝐹_𝑂𝑏𝑗_ + 𝛿𝐹_𝑂𝑏𝑗_) = (𝑀^𝑇^𝑀)^−1^𝑀^𝑇^(𝐹_𝐶𝐹𝐼𝑚𝑔_ + 𝛿𝐹_𝐶𝐹𝐼𝑚𝑔_) and 𝐹_𝑂𝑏𝑗_ = (𝑀^𝑇^𝑀)^−1^𝑀^𝑇^𝐹_𝐶𝐹𝐼𝑚𝑔_.

In the calculation of 𝐴𝐹_𝑁𝑜𝑖𝑠𝑒_, all parameters are set to closely simulate the linear MLS-SIM experimental conditions. We find that collecting 10 different confocal images under 10 different groups of effective PSFs (Figure S4A) is enough to illustrate the working principle. Similar to our experimental implementations, we reconstructed three layers. Since excitation pattern and detection PSF are symmetric with respect to the focal plane, only two layers were effectively reconstructed in the simulation. One is at the focal plane (𝑧_0_ = 0), the other can be anywhere away from the focal plane. We set 𝑧_1_ = ±150 𝑛𝑚 and 𝑧_1_ = ±450 𝑛𝑚 in two different configurations to show its role in multi-layer reconstruction.

In the first configuration (𝑧_1_ = ±150 𝑛𝑚), the effective PSFs from two different layers share very similar patterns. In this case, the columns of 𝑀 in equation E8 are not independent enough to each other to yield reliable solutions by equation E9. Indeed, the noise amplification factor 𝐴𝐹_𝑁𝑜𝑖𝑠𝑒_ is well above unity across the whole spatial frequency space indicating failure of separating two different layers. This result agrees well with the intuition that two layers with separation smaller than the axial resolution limit cannot be properly reconstructed independently.

In the second configuration (𝑧_1_ = ±450 𝑛𝑚), the diversity of effective PSFs between two layers is greatly increased. Due to this reason, two layers can be reconstructed independently over a large range in spatial frequency space (Figure S4B). In comparison to the second layer, the first layer at the focal plane (𝑧_0_ = 0) can be more reliably restored by equation E9 at higher spatial frequencies. This is because the optical transfer functions (OTFs), which are the Fourier transforms of effective PSFs, of the second layer have low passing rate at high spatial frequencies (Figure S4A) and leaves little information to be recovered from noise. Therefore, the proposed multi-layer deconvolution strategy is not to accurately reconstruct multiple layers simultaneously, but to restore a single layer at the focal plane with best possible confinement in axial direction, termed as super-resolution optical sectioning.

The above discussion shows that MLS-SIM preserves independent information from two layers at the same time and these layers can be properly reconstructed by solving a set of linear equations if the confocal images 𝐶𝐹𝐼𝑚𝑔_𝐷𝑘_(𝑥, 𝑦) captured under different groups of effective PSFs are fully sampled. Our implementation of MLS-SIM differs from this discussion in two ways: (1) Multiplicative Richardson-Lucy deconvolution, which is nonlinear in nature, is employed for reconstruction. (2) The captured confocal images 𝐶𝐹𝐼𝑚𝑔_𝐷𝑘_(𝑥, 𝑦) are not fully sampled. Comparing to linear reconstruction, Richardson-Lucy deconvolution naturally applies non-negative regularization and can converge to better solutions than linear deconvolution in many situations. This non-negative constrain, together with the sparsity of the sample, may compensate for missing information from the unsampled pixels in confocal images 𝐶𝐹𝐼𝑚𝑔_𝐷𝑘_(𝑥, 𝑦). In practice, we found multi-layer reconstruction strategy was every effective in cleaning out background interference and promoting reconstruction confinement in axial direction (Figure S4C).

### Phase calibration

The phase calibration established a precise relationship between the measurement from phase grating monitoring photodetector (Figure S1B) and the position of the second structured line illumination pattern on the camera. There were two steps in the calibration. (1) Measure the actual pattern of the second structured line illumination on the camera. Since the circularly distributed grating was employed for phase modulation along the excitation laser line, the period of intensity modulation was not the same along the line (Figure S1C). To calibrate this variation, images of a thin layer of densely packed 200 nm diameter fluorescent beads (F8887 and F8888, Invitrogen) were collected in MLS-SIM while keeping the phase grating static. The captured raw frames were averaged to produce an image of second structured line illumination pattern. The periods of the intensity modulation at different positions on the camera could be determined by measuring the distances between fluorescence peaks in the image. (2) Build the relationship between photodetector’s measurement and the calibrated pattern of the second structured line illumination pattern. To do so, images of a thin layer of densely packed 200 nm diameter fluorescent beads were collected again in MLS-SIM while keeping the phase grating rotating. The phase shift of the excitation pattern captured on each frame was estimated by its cross-correlation with the reference excitation pattern measured in step (1). The calibrated relationship between photodetector’s measurements and estimated phases from many frames could be determined by regression. Based on this calibration, the category of the effective PSF for each pixel could be determined based on the photodetector’s measurement.

### Measurement of excitation and detection PSFs

The measurement of excitation and detection PSFs were performed simultaneously by imaging isolated 100 nm diameter fluorescence bead on glass slide. To measure excitation PSFs of two structured line illumination patterns, the piezo scanning mirror (SM in Figure S1A), which only scanned in y direction in MLS-SIM imaging, performed 2D x-y scan to position the excitation patterns at different positions relative to the isolated fluorescence bead. At each scanning position, an image of the isolated fluorescent bead was collected. Averaging all collected images yielded an excitation PSF with reduced noise. To obtain excitation PSF, all pixel values in each image were summed to estimate the intensity of the excitation pattern experienced by the fluorescent bead at the current scanning position. A 2D image of the excitation pattern could then be generated from this 2D scan. Fluorescence bead was also scanned in the z direction to extend excitation pattern measurement from 2D to 3D. To further minimize measurement noise, phase retrieval was performed using Gerchberg-Saxton algorithm to estimate the detection PSF with the constrain that the intensity of the collected fluorescence was uniform on the objective’s pupil plane (Figure S1E). Phase retrieval was also performed to estimate excitation PSFs using constrains that the excitation laser formed a line in the first excitation pattern and three lines in the second excitation pattern on objective’s back pupil (Figure S1E). To determine nonlinear excitation PSFs, linear excitation PSFs were first measured in the same way described above. They were then updated by introducing nonlinear fluorescence saturation effect according to a model describing the fluorescence saturation effect under pulsed laser excitation^25^ (Figures 6A and 6B). The laser pulse width is 600 ps, and the fluorescence life time was estimated as 3.2 ns (from Gross et al.^61^). The fluorescence emission rate was estimated experimentally by comparing the in-focus signal and background under different excitation laser powers (Figure 6B).

### Estimation of fluorescence emission rate under saturated fluorescence excitations

The nonlinear saturation level could be estimated from varying signal-to-background ratio (SBR) at different excitation laser pulse energies. Due to the saturation effect, the in-focus signal increased nonlinearly with increasing pulse energies, while the background fluorescence, which was excited at much reduced laser intensities away from the focus, increased nearly linearly. To estimate the saturation level, which could be described by nonlinear dependence of fluorescence emission rate on laser pulse energy, a series of images were captured under excitations of different laser pulse energies on the same field of view. For each pixel, the change of pixel values 𝑓(𝑝) with respect to increasing laser pulse energy 𝑝 can be obtained. 𝑓(𝑝) could be considered as a sum of both linear component caused by background fluorescence and nonlinear component from in-focus or near-focus signals: 𝑓(𝑝) = 𝛼 · 𝑝 + 𝛽_1_ · 𝑠_1_(𝑝) + 𝛽_2_ · 𝑠_2_(𝑝) + 𝛽_3_ · 𝑠_3_(𝑝) + ⋯, where the linear component 𝛼 · 𝑝 was caused by background fluorescence and 𝛽_𝑘_ · 𝑠_𝑘_(𝑝) were nonlinear fluorescence emission rate functions caused by in-focus or near-focus signals experiencing different levels of saturations. To simplify the discussion, we searched for 500 pixels in the image that exhibited highest level of nonlinearities in 𝑓(𝑝) and assume that they represented the highest achievable saturation level under the presented excitation pattern and shared the same nonlinear function 𝑠(𝑝). In this case, these 500 pixels shared a common model 𝑓(𝑝) = 𝛼 · 𝑝 + 𝛽 · 𝑠(𝑝) with different coefficients 𝛼 and 𝛽. 𝑠(𝑝) was a parameterized function describing the saturation effect^25^. Using least square fitting, coefficients of 𝛼 and 𝛽 for each pixel and the shared nonlinear function 𝑠(𝑝) could be estimated simultaneously. Given fluorescence emission rate function 𝑠(𝑝) calibrated at experimentally measured laser pulse energies, the nonlinear excitation PSFs could be calculated based on experimentally measured excitation PSFs under linear excitations.

### Zebrafish preparation and imaging

For specifically visualizing the morphology of purkinje cells in zebrafish cerebellum, we generated a construct to drive a codon-optimized sfGFP with a 3’ membrane tag encoding the CAAX box of human Harvey Ras (CTGAACCCTCCTG-ATGAGAGTGGCCCCGGCTGCATGAGCTGCAAGTGTGTGCTCTCC) under the control of a promotor composed of two tandem *cpce* (ca8 promoter derived purkinje cell specific enhancer element) fused to E1B. The *cpce* sequence was PCR-amplified from the zebrafish genome DNA^62^, and the *2×cpce-E1B:sfGFP* was flanked by a minimal *Tol2* transposase recognition sites^63^. The labeling of purkinje cells was achieved via transient transgene by microinjecting the expression plasmid and *Tol2* transposase mRNA into one-cell-stage embryos. The injected plasmid amount was adjusted (here at 15 pg) to result in a moderate mosaic pattern of PC labeling. Larvae were paralyzed with pancuronium dibromide (1 mM), embedded in 1.6% low-melting agarose for volumetric super-resolution imaging using a water immersion objective.

### Mice care and imaging

To image neurons in awake mice, cranial glass windows were implanted before imaging. Mice were anesthetized by isoflurane during the surgery (3% for induction, 1–1.5% during the surgery) and fixed onto the stereotaxic apparatus once fully sedated. Eyes were covered with erythromycin ointment to prevent drying and glare. The scalp was then removed, and the remnants on the skull were removed by scalpel and scissors. The primary somatosensory cortex (S1) was labeled by stereotactic technique (anterior-posterior, -1.0 mm; medial-lateral, 3.8 mm from the Bregma point). A craniotomy of about 3 mm in diameter was administrated with a skull drill above the right hemisphere S1. The craniotomy region was submerged in flowing sterile saline during drilling to cool down the brain surface. After removing the skull, blood and skull debris were carefully washed away with sterile gelatin sponges. The virus mixture was injected with a glass pipette perpendicular to the brain surface with a microinjection syringe pump (UMP3, World Precision Instruments) at a speed of 15 nL/min and with a volume of 100 nL per injection site. Pipette was held for 10 minutes after each injection was completed. After injection, a hat-shaped cranial window, which was made of a circular cover glass (3.2 mm in diameter and 0.17mm thickness) concentrically glued to a cover glass ring (3.0 mm inner diameter, 5.0 mm outer diameter and 0.1mm thickness), was attached to and secured on the skull with dental acrylic. In addition to craniotomy, EEG and EMG electrodes were implanted on mice for imaging experiments during sleep states. For EEG recording, three stainless-steel screws were inserted into the skull on top of the frontal cortex, the S1, and the cerebellum, respectively. For EMG recording, two electrodes were inserted into the neck musculature. All electrodes were secured by dental acrylic. Finally, a head bar was stuck to the skull with glue (Loctite 495) and secured with dental acrylic. Mice were allowed to recover for at least two weeks before imaging.

Before imaging experiments, animals were habituated on the imaging stage every day and successively for one week. To prepare for imaging, mice were head-fixed onto a customized head bar holder without anesthesia. To minimize optical aberrations^23^, the angle of the cranial window was adjusted by a tip-tilt stage such that the surface of the glass window was perpendicular to the optical axis of the imaging objective. During the imaging, mice were freely running on a wheel, or lying on a flat plate in sleep experiments. In sleep experiments, an additional two-hour habituation before imaging was given to mice under the microscope. The acquisition parameters for imaging experiments were summarized in Table S1.

### Image reconstruction and registration

The principle and algorithm for imaging reconstruction were summarized in Figure S2. To prepare for the reconstruction, the actual phase of the second structured line illumination pattern for each captured camera frame was recovered based on the measurement of the phase grating monitoring photodetector (Figure S1B) and its calibrated relationship with the phase of each camera pixel. Based on the recovered phase, raw images captured during a single scan along y axis were re-arranged into 52 confocal images corresponding to 52 groups of effective PSFs. For multi-layer reconstruction, each group consisted of 3 effective PSFs measured at 𝑧 = 0, 𝑧 = 450 nm and 𝑧 = −450 nm from the focal plane. For volumetric imaging that collected images at different depths, the image reconstruction at each depth was independent from each other. In the nonlinear MLS-SIM, raw images were rearranged into 60 confocal images corresponding to 60 groups of effective PSFs. Multi-layer-reconstruction was also employed for image reconstruction in nonlinear MLS-SIM.

MLS-SIM preserved local super-resolution resolving capability, but still suffered from image distortion in presence of strong motions. This image distortion could be corrected most of the time if a time series of 3D volumes were collected. First, the entire time series were manually inspected to remove time points that were too strongly distorted to be recovered. A reference volume was then generated by averaging the remaining time series images. Using this reference volume, the shifts of each volume at each time point were estimated by 3D cross-correlation. Correcting shifts of each time point and averaging them together generated an updated reference volume. This updated reference volume was usually good enough for following row-by-row registration (Figure 2B). Since the MLS-SIM collected a row at a time, the distortion could be described by rigid translation of individual rows in 3 directions. The shifts of each row in the image were found by exhaustive search of best match between a sub-image and the updated reference volume using 2D cross-correlation. For each target row, its sub-image contained N neighboring rows both above and below the target row. When the sample motion was mild, more rows (e.g. N=2 or 3) could be included in the sub-image to improve the registration precision of the target row. This step generated noisy estimations of shifts for each row. These estimations were further improved by curve fitting using a collection of gaussian kernels because the sample motion could be assumed to be continuous. Distorted images were then re-sampled on the pixel grid defined by the reference volume using linear interpolation. Example MATLAB scripts for image registration were included in the online repository.

### Sleep recording and analysis

EEG and EMG electrodes were connected to flexible recording cables via a mini-connector. Recordings started right after mice being placed under the imaging objective. EEG and EMG signals were recorded with a Pinnacle 3-channel system (8200-K1-SL, Pinnacle Technology), high-pass filtered (EEG>0.5 Hz, EMG>10 Hz) and digitized at 1,000 Hz. For sleep scoring, spectral analysis was carried out using fast Fourier transform, and brain states were classified into awake (low delta (1–4.5 Hz) power, low theta-to-delta ratio (6–10 Hz)), NREM sleep (synchronized EEG with high delta power and low EMG power), and REM sleep (high theta-to-delta ratio, low EMG power), with an epoch length of 2.5 s. The classification was made using a custom-written graphical user interface^64^.

### QUANTIFICATION AND STATISTICAL ANALYSIS

#### Quantifications on imaging resolution in presence of sample motions

The resolution limit of the system was quantified by imaging 100 nm diameter fluorescent beads sparsely spread on a glass slide. The same field-of-view was repeatedly imaged under the conditions when the sample was kept static or moved at three different speeds (25, 50, or 100 μm/s) and along eight different directions (0, 45, 90, 135, 180, 225, 270, or 315°). The interspaces between identified bead pairs were measured by fitting a two-peak Gaussian function to their reconstructed images captured under static conditions. When artificial sample motions were introduced, reconstructed images of bead pairs could be distorted but still resolved if separating peaks were preserved. As resolution can be differentially affected by sample motions in different directions, the estimated resolutions across different directions were fitted by cubic spline with MATLAB function *csape* on the interspaces of resolvable bead pairs with different orientations. In each fitting, cubic spline with 4 knots (0, 45, 90, and 135°) and periodic ending conditions was fitted over 20 data points.

#### Quantifications on neural morphologies

Quantifications were performed with custom-written MATLAB scripts and with the help of ImageJ-Fiji^65^ to manually identify regions of interest (ROIs) that exhibited observable morphological features or dynamics. A dendritic spinule was identified as a thin protrusion on a spine with protruding and retracting dynamics during the time-lapse imaging. Spinule exploration ranges were measured as the maximum Euclidean distances between spinules’ roots and their tips during their lifespans. A recurrent spinule site was defined as a location, within 200 nm of which spinules emerge multiple times. The bouton area was measured by manually labeling the region of a continuous swelling on an axon. Further protrusions on axonal boutons were identified with similar criterions as spinules on dendritic spines. For imaging experiments during sleep-wake cycles, the morphologies of boutons, dendrites, and spines were all labeled manually. To quantify the morphological changes on both protrusions and retractions, the alignment between target structures at different time points were optimized to maximize overlaps. The morphology change was defined as the sum of absolute values of both protrusion and retraction. In two-color imaging of PSD-95 and neural morphologies, small protrusions were identified as distinct ridges that introduced angle changes of more than 30 degrees within 500 nm on trunk’s surface (Figure 5E). A trunk PSD near a filopodium or spine was defined as within 500 nm of the root of the filopodium or spine. PSDs were manually classified as small (smaller than 500 nm in all directions), elongated (line-shaped, long axis is longer than 500 nm), or complex-shaped.

The statistical method and P value in each analysis were indicated in the figure legends. The numbers of analyzed samples (n) and standard deviations (SDs) are indicated in the figures or figure legends.

## SUPPLEMENTARY ITEM TITLES AND LEGENDS

**Table S1. Acquisition parameters for MLS-SIM imaging in zebrafish and mice, related to STAR Methods.**

**Video S1. Purkinje cells expressing membrane-targeted EGFP in living larval zebrafish imaged by MLS-SIM, related to Figure 1C.**

Inset shows a magnified view.

**Video S2. Excitatory neurons expressing membrane-targeted EGFP in awake mouse brain. LC, raw line confocal imaging, related to Figure 1D.**

LCDeconv, line confocal imaging with deconvolution. Left, overview of the entire field of view; Right, magnified views of representative ROIs processed as LC (first column), LCDeconv (second column) and MLS-SIM (third column). Moving bars in white boxes indicate the imaging depth of current views.

**Video S3. Motion-distorted images captured in the brain of a head-fixed mouse running on a wheel and correction of motions by image registrations, related to Figure 2.**

MIP, max-intensity projection. Fine structures under severe distortions could be resolved and tracked over time after motion correction.

**Video S4. Time-lapse volumetric imaging in awake mouse brain, related to Figure S3.** Bottom, magnified views. Imaging volumes, consisting of 20 planes, were continuously captures at 5.0 frames/s and over 20 minutes.

**Video S5. Time-lapse volumetric imaging of spinules in awake mouse brain, related to Figure 3.**

Bottom, magnified views of spines with identified spinule emergence. Blink was caused by failures in image registration caused by severe animal motion.

**Video S6. Time-lapse volumetric imaging in mice brain during sleep-wake cycles, related to** **Figure 4**.

Top, brain states inferred from EEG and EMG recording. Gray, wake; yellow, NREM sleep; blue, REM sleep. Dark red bar shows the recording time of the current imaging volume.

**Video S7. Two-color imaging of PSD-95 and membrane of neurons in awake mice brain, related to** **Figure 5**.

Gray, membrane-targeted mRuby3; Red, PSD95.FingR-EGFP. Right, magnified view of representative structures in the volume.

## REFERENCES

1. Debanne, D. (2004). Information processing in the axon. Nat. Rev. Neurosci. 5, 304–316. 10.1038/nrn1397.

2. Scott, E.K., and Luo, L. (2001). How do dendrites take their shape? Nat. Neurosci. 4, 359–365. 10.1038/86006.

3. Holtmaat, A., and Svoboda, K. (2009). Experience-dependent structural synaptic plasticity in the mammalian brain. Nat. Rev. Neurosci. 10, 647–658. 10.1038/nrn2699.

4. Fu, M., and Zuo, Y. (2011). Experience-dependent structural plasticity in the cortex. Trends Neurosci. 34, 177–187. 10.1016/j.tins.2011.02.001.

5. Wefelmeyer, W., Puhl, C.J., and Burrone, J. (2016). Homeostatic Plasticity of Subcellular Neuronal Structures: From Inputs to Outputs. Trends Neurosci. 39, 656–667. 10.1016/j.tins.2016.08.004.

6. White, J.G., Soughgate, E., Thomson, J.N., and Brenner, S. (1986). The structure of the nervous system of the nematode *Caenorhabditis elegans*. Philos. Trans. R. Soc. Lond. B Biol. Sci. 314, 1–340. 10.1098/rstb.1986.0056.

7. Lendvai, B., Stern, E.A., Chen, B., and Svoboda, K. (2000). Experience-dependent plasticity of dendritic spines in the developing rat barrel cortex in vivo. Nature 404, 876–881. 10.1038/35009107.

8. Bock, D.D., Lee, W.-C.A., Kerlin, A.M., Andermann, M.L., Hood, G., Wetzel, A.W., Yurgenson, S., Soucy, E.R., Kim, H.S., and Reid, R.C. (2011). Network anatomy and in vivo physiology of visual cortical neurons. Nature 471, 177–182. 10.1038/nature09802.

9. Nägerl, U.V., Willig, K.I., Hein, B., Hell, S.W., and Bonhoeffer, T. (2008). Live-cell imaging of dendritic spines by STED microscopy. Proc. Natl. Acad. Sci. 105, 18982–18987. 10.1073/pnas.0810028105.

10. Berning, S., Willig, K.I., Steffens, H., Dibaj, P., and Hell, S.W. (2012). Nanoscopy in a Living Mouse Brain. Science 335, 551–551. 10.1126/science.1215369.

11. Tønnesen, J., Katona, G., Rózsa, B., and Nägerl, U.V. (2014). Spine neck plasticity regulates compartmentalization of synapses. Nat. Neurosci. 17, 678– 685. 10.1038/nn.3682.

12. Tønnesen, J., Inavalli, V.V.G.K., and Nägerl, U.V. (2018). Super-Resolution Imaging of the Extracellular Space in Living Brain Tissue. Cell 172, 1108–1121.e15. 10.1016/j.cell.2018.02.007.

13. Arizono, M., Idziak, A., Quici, F., and Nägerl, U.V. (2023). Getting sharper: the brain under the spotlight of super-resolution microscopy. Trends Cell Biol. 33, 148–161. 10.1016/j.tcb.2022.06.011.

14. Sigal, Y.M., Speer, C.M., Babcock, H.P., and Zhuang, X. (2015). Mapping Synaptic Input Fields of Neurons with Super-Resolution Imaging. Cell 163, 493–505. 10.1016/j.cell.2015.08.033.

15. Jacquemet, G., Carisey, A.F., Hamidi, H., Henriques, R., and Leterrier, C. (2020). The cell biologist’s guide to super-resolution microscopy. J. Cell Sci. 133, jcs240713. 10.1242/jcs.240713.

16. Betzig, E., Patterson, G.H., Sougrat, R., Lindwasser, O.W., Olenych, S., Bonifacino, J.S., Davidson, M.W., Lippincott-Schwartz, J., and Hess, H.F. (2006). Imaging Intracellular Fluorescent Proteins at Nanometer Resolution. Science 313, 1642–1645. 10.1126/science.1127344.

17. Xu, K., Zhong, G., and Zhuang, X. (2013). Actin, Spectrin, and Associated Proteins Form a Periodic Cytoskeletal Structure in Axons. Science 339, 452–456. 10.1126/science.1232251.

18. Wu, Y., and Shroff, H. (2018). Faster, sharper, and deeper: structured illumination microscopy for biological imaging. Nat. Methods 15, 1011–1019. 10.1038/s41592-018-0211-z.

19. Masch, J.-M., Steffens, H., Fischer, J., Engelhardt, J., Hubrich, J., Keller-Findeisen, J., D’Este, E., Urban, N.T., Grant, S.G.N., Sahl, S.J., et al. (2018). Robust nanoscopy of a synaptic protein in living mice by organic-fluorophore labeling. Proc. Natl. Acad. Sci. 115, E8047–E8056. 10.1073/pnas.1807104115.

20. Wegner, W., Mott, A.C., Grant, S.G.N., Steffens, H., and Willig, K.I. (2018). In vivo STED microscopy visualizes PSD95 sub-structures and morphological changes over several hours in the mouse visual cortex. Sci. Rep. 8, 219. 10.1038/s41598-017-18640-z.

21. Wegner, W., Steffens, H., Gregor, C., Wolf, F., and Willig, K.I. (2022). Environmental enrichment enhances patterning and remodeling of synaptic nanoarchitecture as revealed by STED nanoscopy. eLife 11, e73603. 10.7554/eLife.73603.

22. Velasco, M.G.M., Zhang, M., Antonello, J., Yuan, P., Allgeyer, E.S., May, D., M’Saad, O., Kidd, P., Barentine, A.E.S., Greco, V., et al. (2021). 3D super-resolution deep-tissue imaging in living mice. Optica 8, 442. 10.1364/OPTICA.416841.

23. Turcotte, R., Liang, Y., Tanimoto, M., Zhang, Q., Li, Z., Koyama, M., Betzig, E., and Ji, N. (2019). Dynamic super-resolution structured illumination imaging in the living brain. Proc. Natl. Acad. Sci. 116, 9586–9591. 10.1073/pnas.1819965116.

24. Heintzmann, R., Jovin, T.M., and Cremer, C. (2002). Saturated patterned excitation microscopy—a concept for optical resolution improvement. J. Opt. Soc. Am. A 19, 1599. 10.1364/JOSAA.19.001599.

25. Gustafsson, M.G.L. (2005). Nonlinear structured-illumination microscopy: Wide-field fluorescence imaging with theoretically unlimited resolution. Proc. Natl. Acad. Sci. 102, 13081–13086. 10.1073/pnas.0406877102.

26. Gustafsson, M.G.L., Shao, L., Carlton, P.M., Wang, C.J.R., Golubovskaya, I.N., Cande, W.Z., Agard, D.A., and Sedat, J.W. (2008). Three-Dimensional Resolution Doubling in Wide-Field Fluorescence Microscopy by Structured Illumination. Biophys. J. 94, 4957–4970. 10.1529/biophysj.107.120345.

27. Mandula, O., Kielhorn, M., Wicker, K., Krampert, G., Kleppe, I., and Heintzmann, R. (2012). Line scan - structured illumination microscopy super-resolution imaging in thick fluorescent samples. Opt. Express 20, 24167–24174. 10.1364/OE.20.024167.

28. Müller, C.B., and Enderlein, J. (2010). Image Scanning Microscopy. Phys. Rev. Lett. 104, 198101. 10.1103/PhysRevLett.104.198101.

29. Sheppard, C.J.R., Mehta, S.B., and Heintzmann, R. (2013). Superresolution by image scanning microscopy using pixel reassignment. Opt. Lett. 38, 2889. 10.1364/OL.38.002889.

30. Mudry, E., Belkebir, K., Girard, J., Savatier, J., Le Moal, E., Nicoletti, C., Allain, M., and Sentenac, A. (2012). Structured illumination microscopy using unknown speckle patterns. Nat. Photonics 6, 312–315. 10.1038/nphoton.2012.83.

31. Schropp, M., Seebacher, C., and Uhl, R. (2017). XL-SIM: Extending Superresolution into Deeper Layers. Photonics 4, 33. 10.3390/photonics4020033.

32. Hering, H., and Sheng, M. (2001). Dentritic spines : structure, dynamics and regulation. Nat. Rev. Neurosci. 2, 880–888. 10.1038/35104061.

33. Kasai, H., Ziv, N.E., Okazaki, H., Yagishita, S., and Toyoizumi, T. (2021). Spine dynamics in the brain, mental disorders and artificial neural networks. Nat. Rev. Neurosci. 22, 407–422. 10.1038/s41583-021-00467-3.

34. Westrum, L.E., and Blackstad, T.W. (1962). An electron microscopic study of the stratum radiatum of the rat hippocampus (regio superior, CA 1) with particular emphasis on synaptology. J. Comp. Neurol. 119, 281–309. 10.1002/cne.901190303.

35. Petralia, R.S., Wang, Y.-X., Mattson, M.P., and Yao, P.J. (2015). Structure, Distribution, and Function of Neuronal/Synaptic Spinules and Related Invaginating Projections. NeuroMolecular Med. 17, 211–240. 10.1007/s12017-015-8358-6.

36. Zaccard, C.R., Shapiro, L., Martin-de-Saavedra, M.D., Pratt, C., Myczek, K., Song, A., Forrest, M.P., and Penzes, P. (2020). Rapid 3D Enhanced Resolution Microscopy Reveals Diversity in Dendritic Spinule Dynamics, Regulation, and Function. Neuron 107, 522–537.e6. 10.1016/j.neuron.2020.04.025.

37. Qiao, Q., Ma, L., Li, W., Tsai, J.-W., Yang, G., and Gan, W.-B. (2016). Long-term stability of axonal boutons in the mouse barrel cortex. Dev. Neurobiol. 76, 252–261. 10.1002/dneu.22311.

38. Tononi, G., and Cirelli, C. (2014). Sleep and the Price of Plasticity: From Synaptic and Cellular Homeostasis to Memory Consolidation and Integration. Neuron 81, 12–34. 10.1016/j.neuron.2013.12.025.

39. Walker, M.P., and Stickgold, R. (2006). Sleep, Memory, and Plasticity. Annu. Rev. Psychol. 57, 139–166. 10.1146/annurev.psych.56.091103.070307.

40. de Vivo, L., Bellesi, M., Marshall, W., Bushong, E.A., Ellisman, M.H., Tononi, G., and Cirelli, C. (2017). Ultrastructural evidence for synaptic scaling across the wake/sleep cycle. Science 355, 507–510. 10.1126/science.aah5982.

41. Yang, G., Lai, C.S.W., Cichon, J., Ma, L., Li, W., and Gan, W.-B. (2014). Sleep promotes branch-specific formation of dendritic spines after learning. Science 344, 1173–1178. 10.1126/science.1249098.

42. Li, W., Ma, L., Yang, G., and Gan, W.-B. (2017). REM sleep selectively prunes and maintains new synapses in development and learning. Nat. Neurosci. 20, 427–437. 10.1038/nn.4479.

43. Maret, S., Faraguna, U., Nelson, A.B., Cirelli, C., and Tononi, G. (2011). Sleep and waking modulate spine turnover in the adolescent mouse cortex. Nat. Neurosci. 14, 1418–1420. 10.1038/nn.2934.

44. Kashiwagi, Y., Higashi, T., Obashi, K., Sato, Y., Komiyama, N.H., Grant, S.G.N., and Okabe, S. (2019). Computational geometry analysis of dendritic spines by structured illumination microscopy. Nat. Commun. 10, 1285. 10.1038/s41467-019-09337-0.

45. Gross, G.G., Junge, J.A., Mora, R.J., Kwon, H.-B., Olson, C.A., Takahashi, T.T., Liman, E.R., Ellis-Davies, G.C.R., McGee, A.W., Sabatini, B.L., et al. (2013). Recombinant Probes for Visualizing Endogenous Synaptic Proteins in Living Neurons. Neuron 78, 971–985. 10.1016/j.neuron.2013.04.017.

46. Marrs, G.S., Green, S.H., and Dailey, M.E. (2001). Rapid formation and remodeling of postsynaptic densities in developing dendrites. Nat. Neurosci. 4, 1006–1013. 10.1038/nn717.

47. Weigert, M., Schmidt, U., Boothe, T., Müller, A., Dibrov, A., Jain, A., Wilhelm, B., Schmidt, D., Broaddus, C., Culley, S., et al. (2018). Content-aware image restoration: pushing the limits of fluorescence microscopy. Nat. Methods 15, 1090–1097. 10.1038/s41592-018-0216-7.

48. Wu, Y., Han, X., Su, Y., Glidewell, M., Daniels, J.S., Liu, J., Sengupta, T., Rey-Suarez, I., Fischer, R., Patel, A., et al. (2021). Multiview confocal super-resolution microscopy. Nature 600, 279–284. 10.1038/s41586-021-04110-0.

49. Hofmann, M., Eggeling, C., Jakobs, S., and Hell, S.W. (2005). Breaking the diffraction barrier in fluorescence microscopy at low light intensities by using reversibly photoswitchable proteins. Proc. Natl. Acad. Sci. 102, 17565–17569. 10.1073/pnas.0506010102.

50. Rego, E.H., Shao, L., Macklin, J.J., Winoto, L., Johansson, G.A., Kamps-Hughes, N., Davidson, M.W., and Gustafsson, M.G.L. (2012). Nonlinear structured-illumination microscopy with a photoswitchable protein reveals cellular structures at 50-nm resolution. Proc. Natl. Acad. Sci. 109, E135–E143. 10.1073/pnas.1107547108.

51. Li, D., Shao, L., Chen, B.-C., Zhang, X., Zhang, M., Moses, B., Milkie, D.E., Beach, J.R., Hammer, J.A., Pasham, M., et al. (2015). Extended-resolution structured illumination imaging of endocytic and cytoskeletal dynamics. Science 349, aab3500. 10.1126/science.aab3500.

52. Hell, S.W., and Wichmann, J. (1994). Breaking the diffraction resolution limit by stimulated emission: stimulated-emission-depletion fluorescence microscopy. Opt. Lett. 19, 780–782. 10.1364/OL.19.000780.

53. Hirano, M., Ando, R., Shimozono, S., Sugiyama, M., Takeda, N., Kurokawa, H., Deguchi, R., Endo, K., Haga, K., Takai-Todaka, R., et al. (2022). A highly photostable and bright green fluorescent protein. Nat. Biotechnol. 40, 1132–1142. 10.1038/s41587-022-01278-2.

54. Liu, Q., Wu, Y., Wang, H., Jia, F., and Xu, F. (2022). Viral Tools for Neural Circuit Tracing. Neurosci. Bull. 38, 1508–1518. 10.1007/s12264-022-00949-z.

55. Qiao, C., Li, D., Guo, Y., Liu, C., Jiang, T., Dai, Q., and Li, D. (2021). Evaluation and development of deep neural networks for image super-resolution in optical microscopy. Nat. Methods 18, 194–202. 10.1038/s41592-020-01048-5.

56. Huang, X., Fan, J., Li, L., Liu, H., Wu, R., Wu, Y., Wei, L., Mao, H., Lal, A., Xi, P., et al. (2018). Fast, long-term, super-resolution imaging with Hessian structured illumination microscopy. Nat. Biotechnol. 36, 451–459. 10.1038/nbt.4115.

57. Chen, Y.-H., Jin, S.-Y., Yang, J.-M., and Gao, T.-M. (2023). The Memory Orchestra: Contribution of Astrocytes. Neurosci. Bull. 39, 409–424. 10.1007/s12264-023-01024-x.

58. Novotny, L., and Hecht, B. (2006). Principles of nano-optics (Cambridge University Press).

59. Lister, J.A., Robertson, C.P., Lepage, T., Johnson, S.L., and Raible, D.W. (1999). nacre encodes a zebrafish microphthalmia-related protein that regulates neural-crest-derived pigment cell fate. Development 126, 3757–3767. 10.1242/dev.126.17.3757.

60. Bensussen, S., Shankar, S., Ching, K.H., Zemel, D., Ta, T.L., Mount, R.A., Shroff, S.N., Gritton, H.J., Fabris, P., Vanbenschoten, H., et al. (2020). A Viral Toolbox of Genetically Encoded Fluorescent Synaptic Tags. iScience 23, 101330. 10.1016/j.isci.2020.101330.

61. Nakabayashi, T., Oshita, S., Sumikawa, R., Sun, F., Kinjo, M., and Ohta, N. (2012). pH dependence of the fluorescence lifetime of enhanced yellow fluorescent protein in solution and cells. J. Photochem. Photobiol. Chem. 235, 65–71. 10.1016/j.jphotochem.2012.02.016.

62. Namikawa, K., Dorigo, A., Zagrebelsky, M., Russo, G., Kirmann, T., Fahr, W., Dübel, S., Korte, M., and Köster, R.W. (2019). Modeling Neurodegenerative Spinocerebellar Ataxia Type 13 in Zebrafish Using a Purkinje Neuron Specific Tunable Coexpression System. J. Neurosci. 39, 3948–3969. 10.1523/JNEUROSCI.1862-18.2019.

63. Balciunas, D., Wangensteen, K.J., Wilber, A., Bell, J., Geurts, A., Sivasubbu, S., Wang, X., Hackett, P.B., Largaespada, D.A., Mclvor, R.S., et al. (2006). Harnessing a High Cargo-Capacity Transposon for Genetic Applications in Vertebrates. PLoS Genet. 2, e169.

64. Barger, Z., Frye, C.G., Liu, D., Dan, Y., and Bouchard, K.E. (2019). Robust, automated sleep scoring by a compact neural network with distributional shift correction. PLOS ONE 14, e0224642. 10.1371/journal.pone.0224642.

65. Schindelin, J., Arganda-Carreras, I., Frise, E., Kaynig, V., Longair, M., Pietzsch, T., Preibisch, S., Rueden, C., Saalfeld, S., Schmid, B., et al. (2012). Fiji: an open-source platform for biological-image analysis. Nat. Methods 9, 676–682. 10.1038/nmeth.2019.

